# Spatial and Temporal Dynamics at an Actively Silicifying Hydrothermal System

**DOI:** 10.1101/2023.02.23.529738

**Authors:** Kalen L. Rasmussen, Blake W. Stamps, Gary F. Vanzin, Shannon M. Ulrich, John R. Spear

## Abstract

Steep Cone Geyser is a unique geothermal feature in Yellowstone National Park, Wyoming (YNP), actively gushing silicon rich fluids along outflow channels possessing living and actively silicifying microbial biomats. To assess the geomicrobial dynamics occurring temporally and spatially at Steep Cone, samples were collected at discrete locations along one of Steep Cone’s outflow channels for both microbial community composition and aqueous geochemistry analysis during field campaigns in 2010, 2018, 2019, and 2020. Geochemical analysis characterized Steep Cone as an oligotrophic, surface boiling, silicious, alkaline-chloride thermal feature with consistent dissolved inorganic carbon and total sulfur concentrations down the outflow channel ranging from 4.59 ± 0.11 to 4.26 ± 0.07 mM and 189.7 ± 7.2 to 204.7 ± 3.55 μM respectively. Furthermore, geochemistry remained relatively stable temporally with consistently detectable analytes displaying a relative standard deviation < 32%. A thermal gradient decrease of ~55°C was observed from the sampled hydrothermal source to the end of the sampled outflow transect (90.34°C ± 3.38 to 35.06°C ± 7.24). The thermal gradient led to temperature-driven divergence and stratification of the microbial community along the outflow channel. The hyperthermophile *Thermocrinis* dominates the hydrothermal source biofilm community, and the thermophiles *Meiothermus* and *Leptococcus* dominate along the outflow before finally giving way to more diverse and even microbial communities at the end of the transect. Beyond the hydrothermal source, phototrophic taxa such as *Leptococcus, Chloroflexus*, and *Chloracidobacterium* act as primary producers for the system supporting heterotrophic growth of taxa such as *Raineya, Tepidimonas*, and *Meiothermus*. Community dynamics illustrate large changes yearly driven by abundance changes of the dominant taxa in the system. Results indicate Steep Cone possesses dynamic outflow microbial communities despite stable geochemistry. These findings improve our understanding of thermal geomicrobiological dynamics and inform how we can interpret the silicified rock record.

## 1 Introduction

Due to the wealth of nutrients and chemical potential creating a plethora of possible ecological niches (Martin et al., 2008; Shock, 1996; Varnes et al., 2003), one of the most popular theories regarding the origin of life revolves around a hydrothermal ecosystem (Pace, 1997; Reysenbach and Cady, 2001; Woese, 1998). Further supporting this theory, some of the oldest phylogenetic lineages are anaerobic thermophiles and hyperthermophiles (Farmer, 2000; Pace, 1997; Reysenbach and Cady, 2001; Setter et al., 1996; Woese, 1998). As a result, hydrothermal environments have become targets for the study of the origin of life and microbial evolution (Varnes et al., 2003). Furthermore, because of the high concentrations of dissolved minerals in the source waters, hydrothermal environments are excellent at lithifying surrounding microorganisms (Walter and Des Marais, 1993). Lithification in hydrothermal systems has proven to be valuable in the preservation and entrainment of microfossils (Walter and Des Marais, 1993) and evidence of past life (Bradley et al., 2017; Kraus et al., 2018; Lowe and Braunstein, 2003). Therefore, hydrothermal lithification processes allow researchers to use current and extinct hydrothermal systems as windows into Earth’s past to inform about ancient life and how life and the surrounding environment have coevolved.

One of the most common types of lithification that occurs in hydrothermal systems is silicification. Silicification of microorganisms is the aggregation of silica particles to cells, resulting in their encrustation within an amorphous silica shell. Silicification is an ideal lithification process for high-resolution microbial preservation due to the mineral’s temporal fidelity (Horodyski et al., 1985). Silica colloids are small, and can preserve cell features such as cell wall pores and even potentially macromolecules (Benning et al., 2005; Schultze-Lam et al., 1996). Due to the recalcitrant nature of precipitated silica, even limited silicification increases the potential for biosignature preservation in hydrothermal systems (Kraus et al., 2018). Because of the excellent morphological preservation, ancient silicified systems are an ideal proxy to better understand the characteristics of both extant and ancient life (Djokic et al., 2017; Schopf, 2006).

The geological interpretations of ancient silicified environments and their entrained microfossils impact how the rock record and the evolution of the geo-biosphere is interpreted, but these geological snapshots may only capture a fraction of their ancient environment. For example, examination of the rock record displays a bias toward the preservation of filamentous and coccoidal morphology (Konhauser et al., 2003; Schopf, 2006), indicating that either other morphologies had not yet evolved or that there exists a preferential preservation bias of certain microorganisms. Additionally, it is known that Mars has deposits of amorphous silica likely from extinct hydrothermal systems (Farmer, 1996; Squyres et al., 2008), and that mineral biosignatures on Earth have been putatively linked to silica deposits on Mars (Ruff and Farmer, 2016). This furthers the belief that extraterrestrial silica deposits are prime locations to search for life and increases the need to understand silicifying hydrothermal systems on Earth.

Previous studies have examined the microbial dynamics in hot spring systems that encompass a wide range of potential variables, their impact on different microbial communities, and have resulted in variable conclusions regarding how dynamic these systems are. Work that examined seasonal impacts on planktonic microbial hot spring communities linked precipitation as a major driver of community structure (Briggs et al., 2014), while an additional study found that precipitation impacts were variable between springs and potentially dependent on aquifer recharge (Colman et al., 2021). Other work found temperature as the greatest selector for microbial communities in sediments and planktonic communities (Guo et al., 2020; Wang et al., 2013), and that planktonic communities were stable temporally while sediment communities were more diverse and dynamic during the sampling period (Wang et al., 2014). Further work has shown that microbiome composition also appears to impact microbial mat communities. For example, biomat biomass and diversity fluctuated during wet and dry seasons in a tropical hot spring (Lacap et al., 2007), while temperate biomat bacteria communities exhibited minimal changes temporally (Ferris and Ward, 1997). The diversity and dynamics of microbial communities that inhabit hot spring environments can impact the preservation of biomarkers. For example, Bosak et al. (2012) showed that in two years, different taxa of cyanobacteria were responsible for the construction of conical microbialites, structures identifiable in the rock record acting as indicators of past life. These studies highlight the irregularity of hot spring environments and highlight the importance of understanding these dynamics to better interpret the rock record, especially in lithifying systems.

One such lithifying system is Steep Cone Geyser (SC; 44.56667 N, 110.8632 W; YNP Research Coordination Network ID: LSMG003), located in the Lower Geyser Basin of Yellowstone National Park (YNP, Figure 1A). Steep Cone has grown vertically from the meadow floor through successive lithification of microbial mats over time and is still rising due to the continued activity of the thermal feature, active silicification, and extensive biomat. The successive lithification at SC has resulted in discrete layers of laminated silicified microbial mats. The lithified structure at SC is amorphous silica in the form of opal-A (Gangidine et al., 2020; Motomura et al., 2003). The various outflow channels at Steep Cone change with time, creating drastic gradients of flourishing microbial communities to fully and recently lithified biomat over a distance of a few meters (Figure 1C, Supplemental Figure S1). Previous research at SC focusing on the silicified sinter deposits found that *Thermus* and *Saccharomonospora* dominated the solid silica deposit at the hydrothermal source, and that microbial cells affected silica deposition and sinter formation (Inagaki et al., 2001). Additional work incorporating SC as one of many thermal features in the study found decreasing sulfide concentrations downstream, which was attributed to microbial sulfide oxidation (Cox et al., 2011). Further research has been conducted that focuses on silica deposition from hydrothermal fluids, particularly the effect of aluminum, and the morphological properties of siliceous deposits at SC (Inagaki et al., 2003; Motomura et al., 2003; Yokoyama et al., 2004). Lastly, potential trace element biosignatures have been assessed in conjunction with fully silicified microbial cells (Gangidine et al., 2020). The goal of this study was to determine the microbial community and geochemical dynamics in the actively silicifying Steep Cone Geyser system to better constrain endpoint conclusions made from examining the ancient, silicified rock record. To address this, geochemistry and biomass samples were collected for simultaneous 16S rRNA and 16S rRNA gene sequencing over multiple years at set locations along an outflow channel of the Steep Cone Geyser.

**Figure 1.**
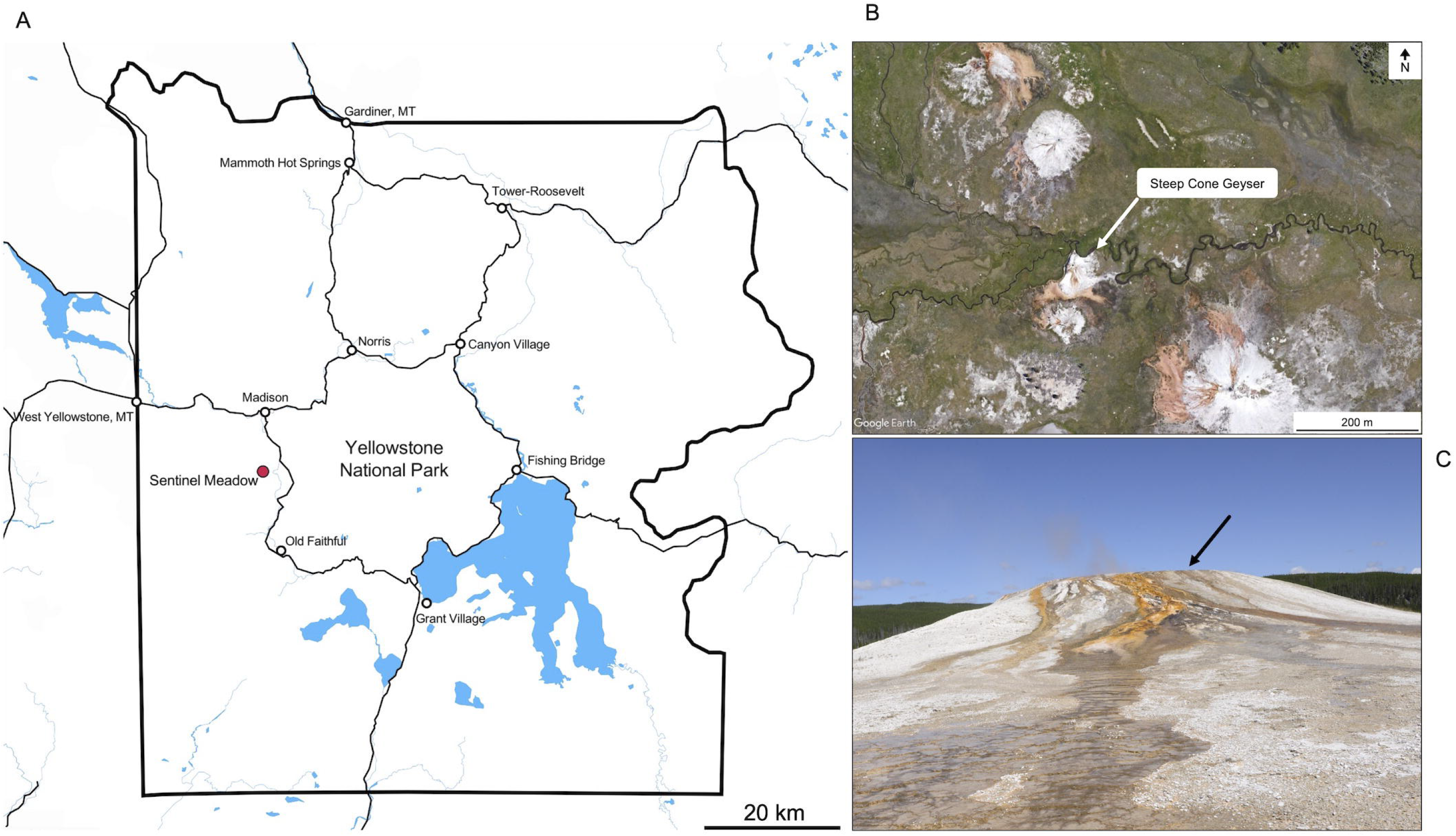
**(A)** Overview of Yellowstone National Park, with Sentinel Meadow demarked by the red dot. Steep Cone Geyser is located within the Sentinel Meadow Hot Springs Group. Map was created in Mapbox (OpenStreetMap contributors, 2022). **(B)** Satellite image of the Sentinel Meadow Hot Springs Group with Steep Cone Geyser indicated by white arrow. **(C)** Image of Steep Cone Geyser from the South taken in August 2021. Black arrow indicates location of sampled outflow channel.

## 2 Materials and Methods

### 2.1 Site Description and Sample Collection

Sample material for this study was collected during seven distinct visits to Steep Cone Geyser in 2010, 2017, 2018, 2019, and 2020 (Supplemental Figure S1). In 2010 (4 June), samples were collected approximately every 1 meter, starting 1 meter from the edge of the vent source, along a vertical transect down the main southeast outflow of the spring to 5 meters. On 18 August 2017, the samples were collected only at the vent source. The vertical transect from 2010 was replicated three times in 2018 (30 May, 26 June, and 29 September), once in 2019 (11 August), and once in 2020 (8 August) plus samples from the vent source (Figure 1C). Samples included: (1) filtered water from the main spring and outflow waters; (2) solid crust/geyserite: (3) swabs of the microbial biofilm; and (4) solid biofilm samples.

Solid surface samples from June 2010 were collected using a sterile spatula, placed in sterile cryovials and stored on ice until they returned to the lab where they were placed in a −20°C freezer until DNA extraction. Before DNA extraction, 2010 samples were homogenized before separation for triplicate DNA extraction. Water filtrate samples collected in 2017, 2018, 2019, and 2020 were filtered through sterilized 0.22 μm hydrophilic polyvinylidene fluoride (PVDF) Durapore^®^ membrane filters using a field washed syringe until clogging occurred (~60 to 1000 ml), and using sterile forceps placed in ZR-96 BashingBead™ lysis tubes containing 750 μl DNA/RNA Shield™ (Zymo Research Corp., Irvine, CA, USA). The samples were shaken and placed on ice until they returned to the vehicle, where they were transferred to a liquid nitrogen dewar. When returning to the laboratory, the samples were stored at −80°C until the extraction of nucleic acid. Triplicate swab samples (2018, 2019, and 2020) were taken along the June 4, 2010 transect of the microbial biomat using sterile BBL™ CultureSwab™ (Becton, Dickinson and Company, Franklin Lakes, NJ, USA), cut with field washed scissors, and stored in the same manner as the water filtrate samples. Fluids for geochemical measurements were filtered using 0.22 μm PVDF Durapore^®^ membrane filter into 50 mL polypropylene tubes for ion chromatography (IC), inductively coupled plasma optical emission spectroscopy (ICP-OES), and inductively coupled plasma atomic emission spectroscopy (ICP-AES) for measurement of major anions and cations. The samples were kept on ice in the field and placed at 4° C upon returning to the laboratory. The samples for ICP-OES and ICP-AES were acidified with 0.2 ml of concentrated nitric acid, shaken, and allowed to equilibrate prior to submission. Samples for dissolved organic and dissolved inorganic carbon (DOC/DIC, 2018, 2019, 2020) were filtered and stored in the same manner in 30 mL combusted amber glass bottles, leaving no headspace for gas exchange. No aqueous geochemistry samples were taken in June 2010, but water samples were collected at the vent source in September 2010 and were diluted 1:25 for ICP analysis as described above. Biomat samples were collected in May 2018 and August 2019 and preserved in 2.5% glutaraldehyde in 1X phosphate buffered saline (PBS) for imaging. The fixed samples were washed three times in 1X PBS and imaged on a Hitachi TM-1000 scanning electron microscope (SEM). A summary of the samples collected, sample types, and collection dates is included in the Supplemental Material (Table S1).

### 2.2 Aqueous Geochemistry

The fluid pH was measured in the field using a rugged Hach Intellical™ PHC101 probe with a portable Hach HQ40d field meter (Hach Company, Loveland, CO, USA). Temperature measurements were collected on site using a Fisherbrand™ Traceable™ noncontact infrared thermometer (Fisher Scientific International, Inc. Hampton, NH, USA). All IC, ICP, and DIC/DOC samples were analysed at the Colorado School of Mines. For 2010 samples, IC and ICP-OES measurements were performed on a Dionex (Thermo Scientific; Waltham, MA, USA) ICS-90 ion chromatography system running an AS14A (4 × 250 mm) column and a Perkin-Elmer (Waltham, MA, USA) Optima 3000 DV ICP-OES system respectively. For fluid samples collected in 2017, 2018, 2019, and 2020, major anions were measured using a Dionex ICS-900 ion chromatography system running an IonPac AS14A-4 μm RFIC (3 × 150 mm) column, while cations were measured using a Perkin Elmer Optima Model 5300 dual view spectrometer ICP-AES. The DOC/DIC measurements were conducted on a Shimadzu total organic carbon analyzer V-TNM-LCSH instrument (Shimadzu Corp. Kyoto, Japan). Geochemical data was analysed in R v4.0.5 (R Core Team, 2022) using the package “Tidyverse” (Wickham et al., 2019) and plots were created using the package “ggplot2” (Wickham, 2016).

### 2.3 DNA/RNA Extractions and 16S rRNA and rRNA Gene Library Sequencing

DNA extraction of the 2010 samples was performed using the MoBio PowerSoil DNA Isolation Kit (MO BIO Laboratories, Inc., Carlsbad, CA, USA), and DNA extraction from the 2017 and 2018 (30 May and 26 July) samples were performed using the ZymoBIOMICS™ DNA MiniPrep kit (Zymo Research Corp.). Simultaneous DNA and RNA extractions were performed on 29 September 2018, 2019, and 2020 samples using the ZymoBIOMICS™ DNA / RNA MiniPrep kit (Zymo Research Corp.). All extractions were performed according to the manufacturer’s instructions, with blank negative extraction controls being used for each round of extractions. Before PCR amplification, RNA samples were converted to cDNA using the SuperScript™ IV First-Strand Synthesis System (Thermo Fisher Scientific Corp., Waltham, MA, USA). DNA contamination in RNA fractions was tested by performing PCR, as outlined below, to check for amplifiable amounts of DNA. If DNA amplification was detected, RNA fractions were diluted until no DNA amplification occurred and diluted samples were used as template for cDNA synthesis. cDNA synthesis was carried out according to the kit manufacturer’s instructions using 1 μL 50 ng/μL random hexamer primers and 2 μL template RNA with parallel positive and negative reverse transcription controls. Synthesized cDNA was directly used as template inputs for subsequent PCR reaction. Archaeal and Bacterial 16S rRNA and rRNA gene libraries were amplified by PCR using primers that flank the V4 region between the 515 and 926 base pair positions *(E. coli* reference). Primers 515F_Y and 926R were chosen for their ability to amplify the V4 region, as described by Parada et al. (2016). The forward primer (M13-515_Y: 5’-**GTA AAA CGA CGG CCA G**<u>TC CGT GYC AGC MGC CGC GGT AA</u>-3’) contains the M13 forward primer (bold) ligated to the 16S rRNA gene-specific sequence (underlined) to facilitate barcoding in a later PCR reaction (Stamps et al., 2016). The utility of the reverse primer (926R: 5’-CCG YCA ATT YMT TTR AGT TT-3’) was illustrated by Parada et al. (2016).

PCR was carried out in 25 μL reactions consisting of 1X 5PRIME HOT MasterMix (Quantabio, Beverly, MA, USA), 0.2 μM of each primer, molecular grade water and 2 μL extracted template DNA. Template DNA was diluted to < 10 ng/μL prior to PCR amplification. The PCR amplification was conducted as follows; initial denaturation at 94°C for 2 min, 30 cycles of (94°C for 45 sec, 50°C for 45 sec, and 68°C for 1:30 min), a final extension of 68°C for 5 min, and a final 4°C hold. The amplicons were cleaned using KAPA Pure Beads (KAPA Biosystems Inc., Wilmington, MA, USA) at a final concentration of 0.8X v/v to remove the solution of primer dimers. Next, a second six-cycle PCR amplification was conducted to add unique 12 basepair barcodes to each cleaned sample amplicon (Hamady et al., 2008) using the forward primer composed of the barcode plus M13 forward sequence (5’-3’) and the 926R primer. The final barcoded PCR amplicons were again cleaned using the KAPA Pure Beads at the above conditions and quantified using the Qubit^®^ dsDNA HS assay (Life Technologies, Carlsbad, CA, USA). Lastly, the amplicons were combined in equimolar amounts, and concentrated using two Amicon^®^ Ultra-0.5 mL 30K Centrifugal Filters (EMD Millipore, Billerica, MA, USA) to a final volume of 80 μL. The final pooled library was submitted for high throughput sequencing on the Illumina MiSeq platform using the PE250 V2 chemistry method (Illumina, San Diego, CA, USA). For this work, all samples were prepared for DNA sequencing in the Geo-Environmental Microbiology Laboratory at the Colorado School of Mines, while sequencing was conducted at the Duke Center for Genomic and Computational Biology (Duke University, Durham, NC, USA).

### 2.4 Amplicon Sequence Processing and Analysis

After sequencing, reads were demultiplexed using AdapterRemoval version 2 (Schubert et al., 2016), and forward and reverse primer sequences were excised using Cutadapt version 3.5 resulting in a final length of 198 bases for the forward reads and 230 bases for the reverse reads (Martin, 2011). DADA2 was used to assess quality (Callahan et al., 2016). All reads possessed a quality score greater than 30 over the length of the trimmed reads for both the forward and reverse sequences and no further trimming was conducted. DADA2 was used to filter samples using standard filter parameters, perform sample inference, merge paired-end Illumina reads, construct an amplicon sequence variant (ASV) table at 100% similarity, remove chimeric sequences, and assign taxonomy using the SILVA database release Version 138 (Pruesse et al., 2007). A phylogenetic tree was constructed using FastTree (Price et al., 2010) for microbial community composition analysis. Contamination reads were identified and removed from samples using Decontam version 1.10 (Davis et al., 2018). Bacteria and Archaea 16S rRNA gene amplicon samples were gene copy number normalized based on the *rrn*DB database version 5.7 as shown previously (Chen et al., 2020; Stoddard et al., 2015).

For within-sample alpha diversity calculations, ASV singletons were removed from the dataset. Then sample richness was estimated using Breakaway, and sample evenness was calculated using the R package Microbiome (Willis and Bunge, 2015; Leo Lahti et al., 2017). The Kruskal-Wallis and pairwise Wilcoxon tests were used to determine alpha diversity statistical significances (adjusted *p*-value < 0.05), along with the Wilcoxon effect size. To quantitatively assess between sample diversity, samples were first bootstrap rarefied with replacement (63 trials) to a sequencing depth of 1080 as previously performed (Salmon et al., 2022). Bray-Curtis Dissimilarity distance matrices were constructed using the R package Vegan, and beta diversity plots were constructing using ggplot2 and loess regression for line fitting (Bray and Curtis, 1957; Oksanen et al., 2022; Wickham, 2016). Permutational analysis of variance (PERMANOVA) and pairwise comparisons (adjusted *p*-value < 0.05) were calculated using Vegan to test statistical significance between beta diversity groupings between dates and locations (10000 permutations). A similarity percentages (SIMPER) analysis was conducted at the genus level while removing any ASVs that contributed less than 0.5% (function simper.pretty in R https://doi.org/10.5281/zenodo.4270481) to determine genera responsible for driving differences in microbial communities (Clarke, 1993). Resultant statistically significant ASVs were determined using the Kruskal-Wallis test for multiple comparisons (adjusted *p*-value < 0.05, function kruskal.pretty in R https://doi.org/10.5281/zenodo.4270481).

ASV ranks where no classification was assigned were fixed using MicroViz to assign distinct identifiers from higher phylogenetic ranks and allow for proper aggregation at lower taxonomic ranks (i.e. genus level) (Barnett et al., 2021). Heatmaps were constructed using Ampvis2 to examine microbial taxa through both time and space, with relative abundances calculated at the genus level (Andersen et al., 2018). ASVs were aggregated at the genus level for differential abundance analysis using the R package Corncob (Martin et al., 2020). Pairwise differential abundance comparisons were made between sequential dates or locations with an adjusted *p*-value < 0.05 cutoff. The most abundant ASVs were compared to other published sequences using the NCBI BLASTN tool to determine their percent similarity to closely related taxa (National Library of Medicine (US), 1988). Commands used to conduct sequence processing, analysis, and figure construction are publicly available at (https://github.com/kalen-rasmussen/Steep-Cone-Temporal-Spatial-Study).

## 3 Results

### 3.1 General Field Observations and Biofilm Descriptions

The visual transformation of the sampled outflow channel at SC over the sampling period is illustrated in Supplementary Figure 1. Down the outflow channel the biomat microbial communities transitioned from opaque microbial communities to pigmented communities dominated by green, browns, yellow, and orange. Pigmented communities began as green/brown biomat but transitioned to yellow and orange biomat over the sampling time course. Upon initial sampling in 2010, the sampled outflow channel was visually one of the main outflow channels at SC with high flow; however, over the sampling period the flow rate along the transect gradually decreased. The transient nature of outflow channels at SC is seemingly important for its accretion overtime as attributed to its height off the meadow floor and mostly uniform cone shape (disregarding where rockfalls have occurred). Depth of fluid flow across the sampled transect decreased over the sampling period from ~1 cm to ~2-5 mm. During sampling, high winds were experienced on the top of SC, often from the east in the morning and from the west in the afternoon. These high winds pushed outflow channel waters laterally, saturating neighboring dried and lithified deposits or forming pools on the flat portions at the top of the feature.

SEM analysis of silica sinter samples at the hydrothermal source matched the descriptions of previous studies at SC (Inagaki et al., 2001; Motomura et al., 2003). In brief, sinter deposits were highly cemented by amorphous silica and silica colloids. Thin actively growing microbial biofilms occurred sporadically on the sinter surface, and entombed cells were visible in the cemented sinter. Samples along the outflow channel were characterized as biofilm ranging from ~1 to 3 mm in thickness. Filamentous cell morphologies bound together by extracellular polymeric substances (EPS) comprised the biofilm. Cells appeared predominantly parallel to the feature surface and randomly oriented in lateral directions across the surface. The EPS and random orientation of cells created a distinct and cohesive biofilm layer sitting atop a fully silicified cell matrix. The underlying silicified matrix was composed of fully silicified microbial filaments and amorphous silica colloid deposits. The silicified matrix exhibited a gradient of cementation and infilling from the hydrothermal source to the transect terminus, with samples at 1 m being highly porous to samples at 5 m exhibiting high degrees of amorphous infilling. See Supplementary Figures 2 and 3 for SEM imagery at the different SC locations.

### 3.2 Geochemical Dynamics

Steep Cone Geyser is generally classified as an oligotrophic, surface boiling, silicious, alkaline-chloride thermal feature. The hydrothermal source waters were sampled seven times throughout the study, and the main parameters are shown in Table 1. The source waters were geochemically stable during the course of the study, especially considering the dynamic nature of YNP and the geothermal features within the park. The mean temperature and pH of the source waters were 90.34°C ± 3.38 and 7.98 ± 0.47 respectively, which is roughly surface boiling at YNP altitude (~2440 m). The mean dissolved inorganic carbon was 4.42 ± 0.25 mM while the dissolved organic carbon ranged from below the limit of quantification (LOQ = 1.42×10^-2^) to 0.7 mM. Nitrate and nitrite concentrations also ranged from below the detection limits (LOQ = 1.61×10^-3^ and 2.17×10^-3^) to 0.93 and 7.4×10^-3^ mM, respectively, with the 0.93 mM nitrate measurement being an outlier. Despite its biological importance, phosphorus sources were mostly below the limit of quantification, with a high total phosphorus measurement of 0.9 μM and a single measurement above the limit of quantification of 9 μM for phosphate. At the source, total sulfur and sulfate were in high abundance compared to other potential metabolic analytes possessing means of 0.23 ± 0.08 and 0.16 ± 0.004 mM respectively. As part of a different study, total sulfide in the source waters was 12.63 ± 1.84 μM on August 8, 2021 (Rasmussen, et al., in preparation). Arsenic has been shown as a potentially important electron donor in hot spring systems (Kulp et al., 2008; Oremland and Stolz, 2003), and total arsenic averaged 14.5 ± 1 μM in the source waters during the sampling period. Lastly, the mean dissolved total silicon in the source waters was 5.28 ± 0.28 mM and chloride averaged 7.32 ± 0.31 mM.

**Table 1.** Aqueous geochemistry values (in mM or °C) from the vent source fluids of Steep Cone Geyser. Entries presented as (BDL) were below the detection limit of the instrument. Entries presented as (NM) were not measured for that geochemical analyte. The LOQ column indicates the limit of quantification for each parameter and methodology (in mM). Subscript (T) indicated total concentration of the given analyte.

| Analyte | 2010-9-23 | 2017-8-18 | 2018-5-30 | 2018-7-26 | 2018-9-29 | 2019-8-11 | 2020-8-21 | LOQ |
| --- | --- | --- | --- | --- | --- | --- | --- | --- |
| Al<sub>T</sub> | 0.0102 | 0.0111 | 0.0106 | 0.0108 | 0.0112 | 0.0106 | 0.0107 | 1.56E-04 |
| As<sub>T</sub> | 0.0138 | 0.0149 | 0.0154 | 0.0151 | 0.0151 | 0.0145 | 0.0126 | 1.07E-04 |
| B<sub>T</sub> | 0.3005 | 0.2789 | 0.2912 | 0.3063 | 0.3035 | 0.2905 | 0.3105 | 2.87E-04 |
| Br<sup>-</sup> | NM | 0.0104 | 0.0082 | 0.0096 | 0.0101 | 0.0140 | 0.0090 | 1.25E-03 |
| Ca<sub>T</sub> | 0.0088 | 0.0053 | 0.0047 | 0.0045 | 0.0055 | 0.0096 | 0.0103 | 1.22E-04 |
| Cl<sup>-</sup> | NM | 7.0759 | 7.0201 | 7.0421 | 7.6061 | 7.6947 | 7.5300 | 2.82E-03 |
| DIC | NM | NM | NM | 4.7316 | 4.4818 | 4.1297 | 4.3661 | 1.42E-02 |
| DOC | NM | NM | NM | BDL | BDL | 0.2600 | 0.6969 | 1.42E-02 |
| F<sup>-</sup> | NM | 1.5501 | 1.4799 | 1.4413 | 1.5543 | 1.7496 | 1.2738 | 5.26E-03 |
| Fe<sub>T</sub> | 1.75E-04 | 1.06E-04 | 2.11E-04 | 5.03E-05 | BDL | BDL | 1.95E-04 | 5.37E-06 |
| Li<sub>T</sub> | 0.2450 | 0.2175 | 0.2410 | 0.2218 | 0.2090 | 0.1984 | 0.2197 | 4.75E-04 |
| Mg<sub>T</sub> | 0.0002 | BDL | BDL | BDL | BDL | 0.0011 | 0.0018 | 1.15E-04 |
| Mn<sub>T</sub> | 2.24E-05 | 1.34E-05 | 1.04E-05 | 1.14E-05 | BDL | 1.28E-05 | 3.28E-05 | 1.82E-06 |
| NO<sub>3</sub><sup>-</sup> | NM | BDL | 0.9308 | BDL | BDL | BDL | BDL | 1.61E-03 |
| NO<sub>2</sub><sup>-</sup> | NM | 0.0074 | BDL | 0.0039 | BDL | BDL | BDL | 2.17E-03 |
| pH | 7.94 | 8.1 | 8.07 | 8.41 | 8.4 | 7 | 7.91 | 0 |
| PO<sub>4</sub><sup>3-</sup> | NM | BDL | 0.0090 | BDL | BDL | BDL | BDL | 1.05E-03 |
| P<sub>T</sub> | BDL | BDL | 0.0009 | BDL | BDL | BDL | 4.16E-04 | 6.46E-05 |
| K<sub>T</sub> | 0.2570 | 0.2493 | 0.2635 | 0.2607 | 0.2569 | 0.2403 | 0.2397 | 4.22E-03 |
| Si<sub>T</sub> | 5.5246 | 4.9959 | 5.3816 | 5.3106 | 5.7156 | 5.1154 | 4.9700 | 1.25E-04 |
| Na<sub>T</sub> | 11.8216 | 9.9967 | 11.3394 | 11.2225 | 11.4372 | 10.7263 | 11.1503 | 1.25E-03 |
| SO<sub>4</sub><sup>2-</sup> | NM | 0.1664 | 0.1566 | 0.1566 | 0.1591 | 0.1570 | 0.1582 | 1.04E-03 |
| S<sub>T</sub> | 0.4066 | 0.1737 | 0.1790 | 0.1899 | 0.2119 | 0.2125 | 0.2086 | 6.17E-04 |
| Temp. | 93.6 | 87.78 | 90.5 | 91.1 | 88.4 | 93 | 88 | - |

Hydrothermal environments can exhibit large spatial geochemical gradients. The outflow channel at SC exhibited a steep thermal gradient, with waters near surface boiling at the source and decreasing to a mean of 35.06 ± 7.24 °C after 5 m, a decrease of ~55 °C (Figure 2). pH displayed a less drastic gradient increasing from an average of 7.98 ± 0.52 in the source waters to 8.28 ± 0.55 5 m down the outflow. Dissolved inorganic and dissolved organic carbon remained stable along the outflow channel ranging from 4.42 ± 0.25 at the source to 4.14 ± 0.13 mM 5 m downstream and 0.24 ± 0.32 to 0.18 ± 0.31 mM, respectively. Similarly, total sulfur concentrations were stable down transect with averages ranging from 190.34 ± 7.34 to 197.43 ± 11.1 μM. Alternatively, mean sulfate concentrations increased from 159 ± 3.76 μM at the thermal source to 191.45 ± 17.43 μM at the transect terminus. Nitrate, nitrite, phosphate, and total phosphorus concentrations remained low along the outflow channel, with most samples occurring below the limit of quantification (Supplemental Figure S4). Mean total arsenic increased from 14.57 ± 1.04 to 16.2 ± 1.43 μM down the sampling transect. Lastly, chloride, a conservative tracer for evaporation, increased from 7.33 ± 3.13 at 0 m to 8.5 ± 4.84 mM at 5 m.

**Figure 2.**
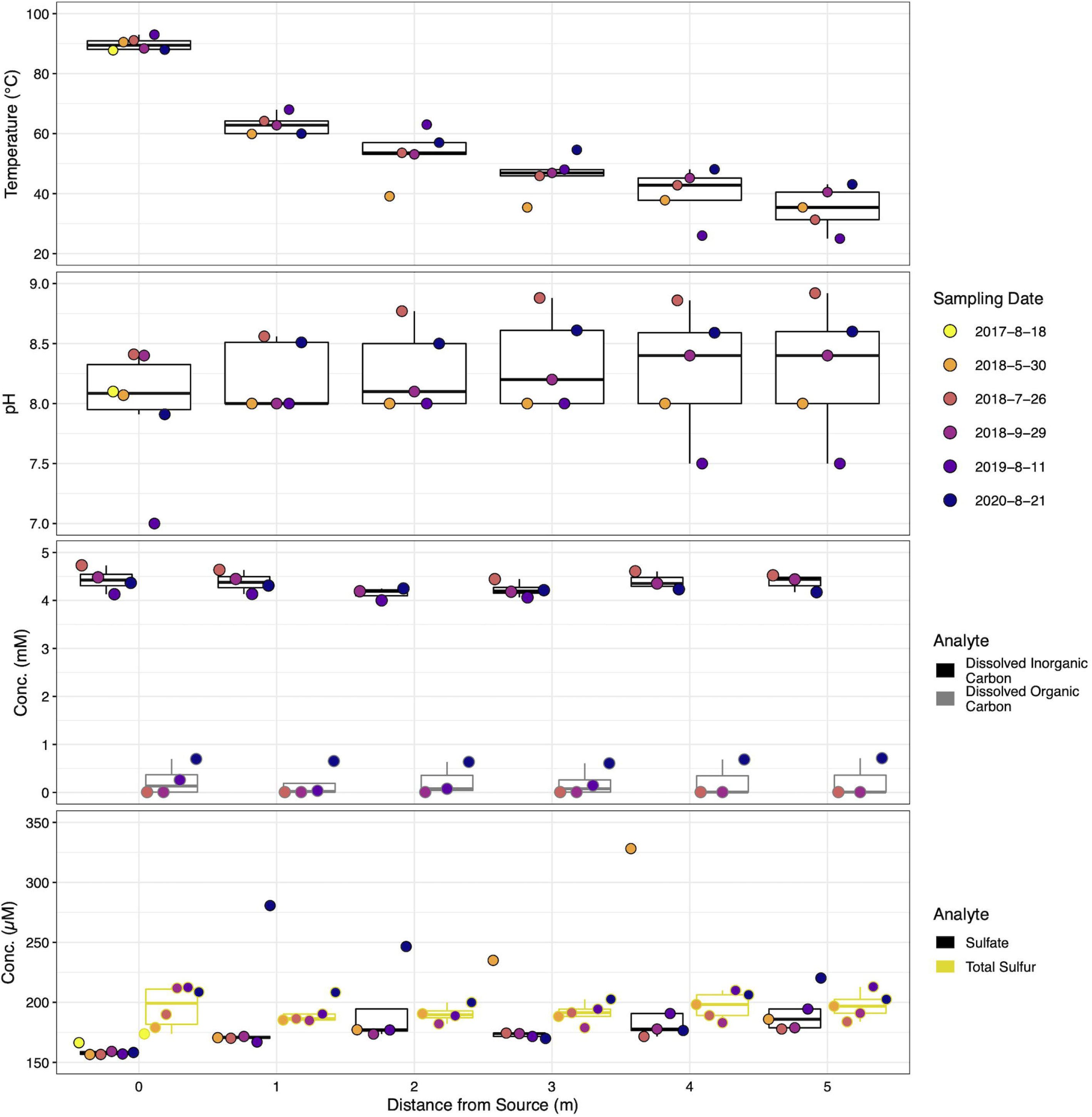
Dot and box plot illustrating geochemical stability down the sampled transect across sampling dates of parameters of interest. Filled dot colors indicate sampling date while box colors indicate measured parameter. Dots are arranged from left to right, oldest to most recent samples for each distance, respectively. Dates are shown as year-month-day.

Although temporal geochemical characterization of the SC source waters indicated a relatively stable influent for the outflow transect, across all locations average geochemical trends were dynamic. The average temperature of all sample locations increased steadily from 49.7 ± 22.05 to 58.5 ± 15.71 °C between May 2018 and August 2020. Mean dissolved organic carbon and total sulfur increased from below the limit of quantification to 0.66 ± 0.06 mM and 189.7 ± 7.2 to 204.7 ± 3.55 μM, respectively. Inversely, mean dissolved inorganic carbon and total arsenic decreased from 4.59 ± 0.11 to 4.26 ± 0.07 mM and 16.9 ± 0.8 to 13.5 ± 0.6 μM respectively over the same period. The trend of decreasing dissolved inorganic carbon and total arsenic occurred temporally at all sample locations. Similarly, increasing organic carbon temporally occurred at all sample locations. The average concentrations of sulfate began high at 208.9 ± 64.25, decreased to 170 ± 8.1 in July 2018, only to rebound back to 208.7 ± 48.6 μM in August 2020. Minimal temporal trends at individual locations were observed for sulfate and total sulfur, while pH measurements were stochastic. Consistently detectable analytes displayed a relative standard deviation < 22% over time across all locations, excluding sulfate (< 32%).

### 3.3 Microbial Community Spatial and Temporal Dynamics

Pielou’s evenness is a metric that quantifies microbial community taxa representation, with one being a completely even community and zero being a community composed of a single species (Pielou, 1966). At SC, evenness was lowest at the vent source and increased down the outflow channel (Figure 3). Evenness began at ~0.5 at the hydrothermal source and plateaued after 2 m between 0.6 and 0.8. Both spatial and temporal effects were statistically significant, *p* = 0.00253 and *p* = 8.46×10^-5^, respectively. Pairwise statistical tests between dates indicated six statistically significant shifts between dates (excluding 2017 due to only sampling at the source). Of note temporally, the decrease in evenness between 2010 and 2020 displayed the greatest effect size (*r* = 0.64, *p* = 0.005) followed by September 2018 to 2020 (*r* = 0.58, *p* = 0.007). Seven spatial pairwise comparisons were statistically significant, with all evenness comparisons to 0 m determined to be significant. The largest effect size was between 0 and 4 m (*r* = 0.59, *p* = 0.019).

**Figure 3.**
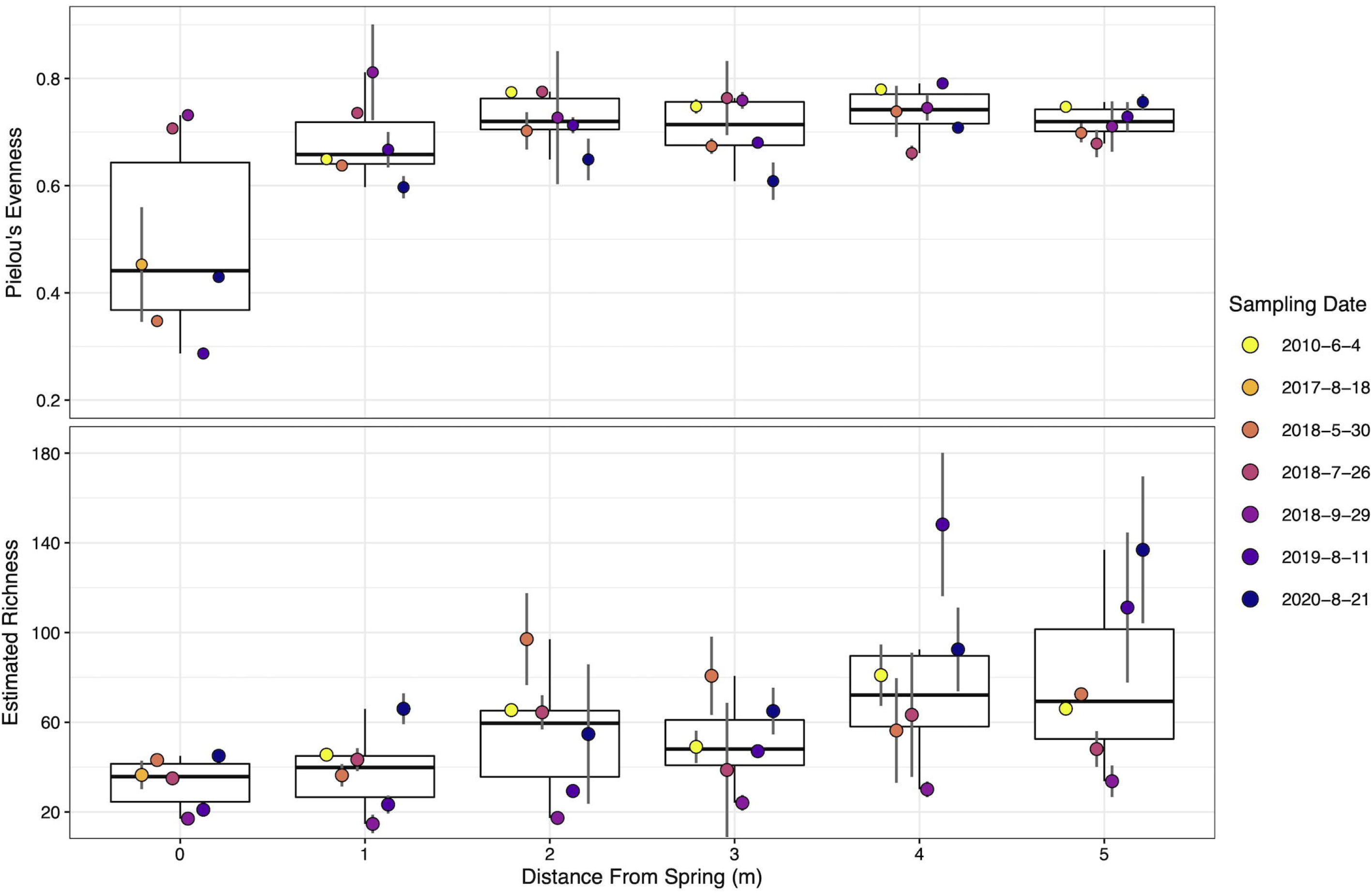
Dot and box plot of alpha diversity metrics down the sampled transect across the different sampling dates. Filled dot colors indicate sampling date. Dots are arranged from left to right, oldest to most recent samples for each distance respectively. Top – Pielou’s evenness. Bottom – Estimated Richness using Breakaway. Error bars indicate the standard deviation calculated from biological replicates (n = 3). Dates are shown as year-month-day.

Estimated richness, a metric estimating the total number of different taxa in the environment from the sample population (Willis and Bunge, 2015), steadily increased down transect, ranging from ~35 taxa at the source to ~80 taxa at the transect terminus. Spatial and temporal effects proved to be statistically significant, *p* = 4.2×10^-4^ and *p* = 8.7×10^-7^, respectively. Pairwise statistical tests found six temporally and four spatially significant changes in richness. The estimated richness in September 2018 was significantly lower than all other sampling dates (excluding 2017 due to sampling), with the largest effect size between September 2018 and 2010 *(r* = 0.84, *p* = 5.8×10^-7^). July 2018 and 2020 richness significantly shifted *(r* = 0.54, *p* = 0.006), a difference especially evident at locations greater than 3 m. Estimated richness increased from 0 m to both 4 m (*p* = 0.035, *r* = 0.51) and 5 m (*p* = 0.009, *r* = 0.59), and 1 m to 4 m (*p* = 0.007, *r* = 0.54) and 5 m (*p* = 0.004, *r* = 0.58). Temporal trends in richness at individual locations appeared stochastic and no statistically significant shifts were determined (Figure 3). Statistical results for pairwise evenness/richness temporal and spatial comparisons are summarized in Table S2.

The between sample diversity was examined both temporally and spatially using the Bray-Curtis dissimilarity metric (Figure 4). Permanova analysis determined location (*R^2^* = 0.06, *p* = 0.025) and time (*R^2^* = 0.22, *p* = 0.0015) as significant. Temporal-spatial interactions were determined to be the most impactful *(R^2^* = 0.28, *p* = 0.0015), indicating significant shifts in community composition occur over both time and space. Overall, two distance comparisons and 12 temporal comparisons displayed significant shifts in the microbial community composition (Table S3). The microbial community became significantly less similar from 2 to 3 m (*R^2^* = 0.064, *p* = 0.03) and 3 to 4 m (*R^2^* = 0.066, *p* = 0.019). All temporal pairwise comparisons with the initial 2010 samples and 2020 samples were determined to be statistically different (Table S3).

**Figure 4.**
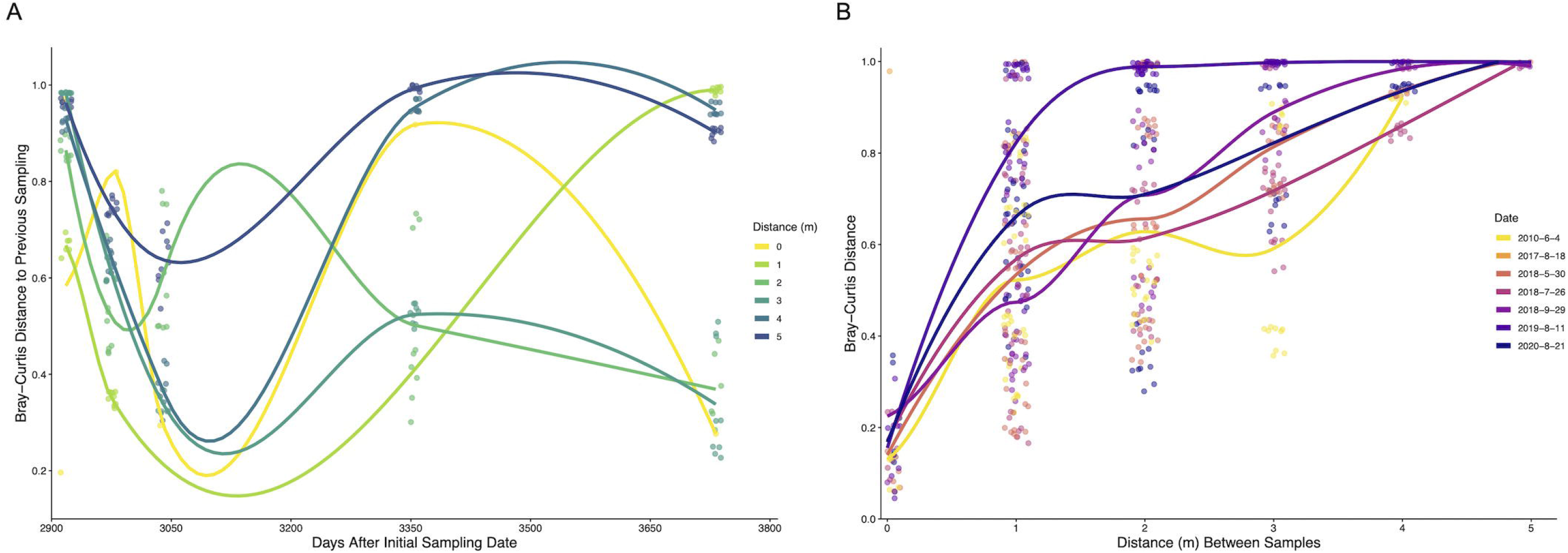
**(A)** Time-lag of Bray-Curtis distances of bacterial and archaeal communities of biofilm samples sampled at Steep Cone Geyser. Dots represent pairwise comparisons between a sample and the previous sampling date. **(B)** Distance-lag plot of Bray-Curtis distances of bacterial and archaeal communities of biofilm samples sampled at Steep Cone Geyser. Dots represent pairwise comparisons between two samples at different distances. Trend lines were determined using Geom_smooth in the R package stats::loess, with trends being fit using loess regression. Dates are shown as year-month-day.

Figure 4A illustrates the pairwise Bray-Curtis dissimilarity between a sample and the corresponding previous sampling date since the initial 2010 sampling. Initial comparisons indicated high dissimilarity across all locations sampled; however, the microbial communities became more similar over time across the first three time points (excluding 0 m), which corresponds to the 2018 sampling dates and less time between sampling events. After 3040 days since initial sampling (pairwise between July and September 2018 samples) the microbial communities across different locations became more stochastic and exhibited increasing variance. Indicating clear shifts in microbial community composition increased in magnitude with increasing time between samplings. Only 2, 3, and 4 m distances possessed significant temporal-spatial shifts. Figure 4B shows the Bray-Curtis dissimilarity between two samples taken at different distances from the hydrothermal source with increasing distance between sampling locations. Microbial communities at the same location, across sampling dates, were highly similar and ranged from ~0.05 to ~0.5, while samples the largest distance apart (5 m) were completely dissimilar (~1). The microbial communities became less similar as the distance between samples increased for all sampling dates. Temporal and spatial variance comparisons between different dates or locations displayed similar distributions (Supplemental Figure S5). The median Bray-Curtis temporal-spatial distance was the highest 0.88, which was significantly more than the temporal (*p* = 3.3×10^-23^) and spatial (*p* = 5.9×10^-12^) variances, indicating that the covariate of location and distance impacts the diversity of the microbial community more than the corresponding individual variates.

### 3.4 Microbial Community Composition and Impact

The relative abundances of the 15 most abundant microbial taxa were examined to elucidate the compositions of the microbial communities and the temporal-spatial changes (Figure 5). The top 15 taxa captured a majority of the total abundance across sampling times and locations ranging from 74% to 100%, excluding 2019 samples at 4 m (30.4%) and 5 m (35.7%). The hyperthermophile genus *Thermocrinis* dominated the hydrothermal source microbial biofilm community, comprising greater than 82% across all sampling dates (Huber et al., 2015), with large contributions (> 12.5%) from *Thermus* ASVs in July and September 2018. Overall, *Meiothermus* dominated the biofilm system and was the most abundant genus across sampling dates and locations. When queried using NCBI BLAST, the most abundant *Meiothermus* ASV was 99% identical to *Meiothermus ruber* (heterotypic synonym *Thermus ruber)*, a red pigmented obligately thermophilic, obligately aerobic heterotroph isolated from a hot spring environment (Loginova et al., 1984). Relative abundances of *Meiothermus* ranged drastically beyond the hydrothermal source, ranging from 0.2% at 5 m in 2019 to 77% at 2 m in September 2018. *Tepidimonas* was the next most abundant putative aerobic heterotrophic microorganism and shared 99% similarity with *Tepidimonas taiwanensis* (Chen et al., 2006). *Tepidimonas* comprised 11.6% of the community at 1 m in 2010 but was mostly absent from locations further down the outflow and July/September 2018 samples. The genera *Raineya* and *Lewinella* round out the remaining putative aerobic heterotrophs in the top 15 taxa (Albuquerque et al., 2018; Khan et al., 2007).

**Figure 5.**
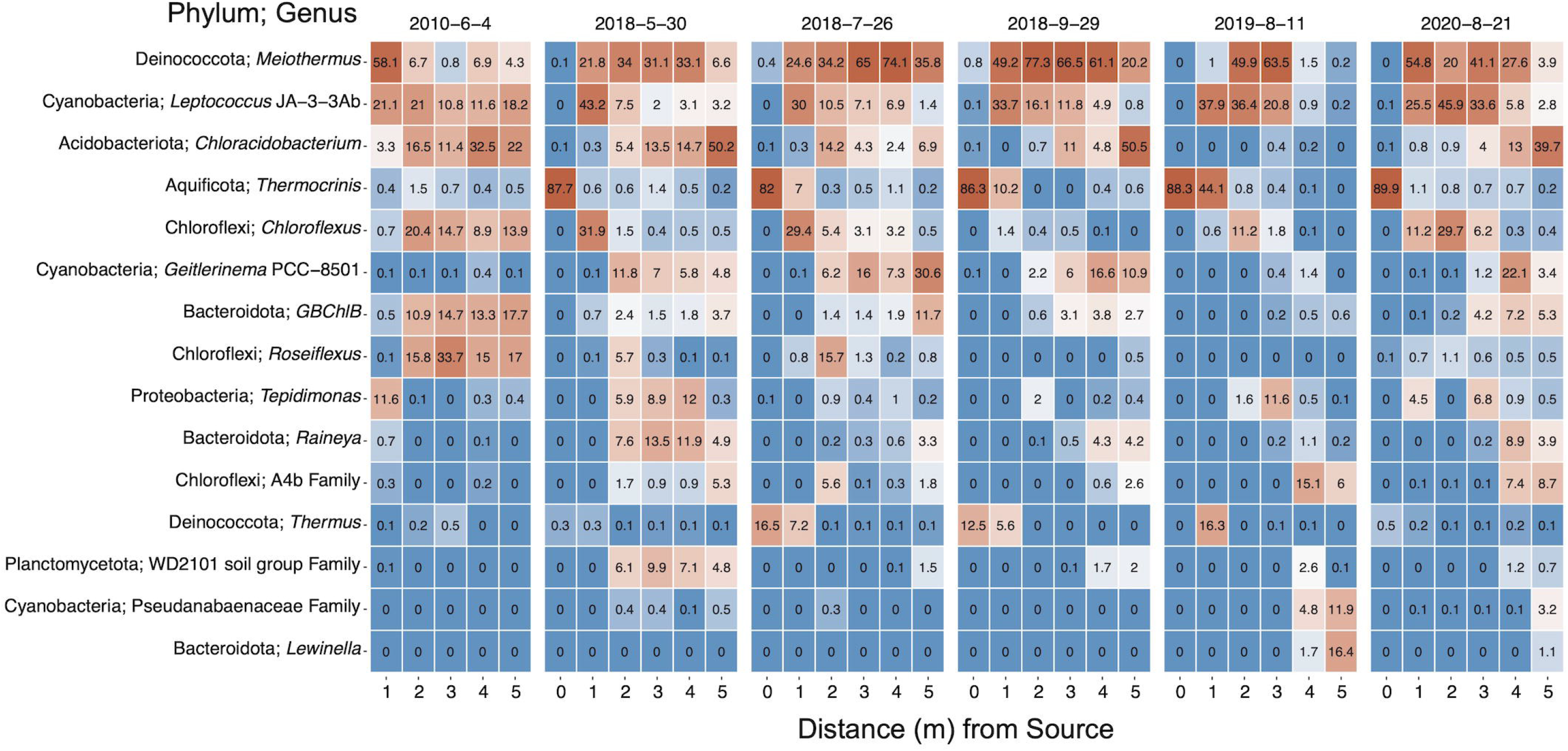
Heat map of the top 15 ASVs within the bacterial and archaeal communities from 16S rRNA gene sequencing. ASVs are named by phyla and most likely genera. Values indicate the mean percent relative abundance. Data is faceted by sampling date and ordered down the sampling transect. Phylum; Genus or lowest classification ID. Dates are shown as year-month-day.

Potential sulfur oxidizers (*Thermocrinis* and *Tepidimonas*), sulfur reducer GBChlB, and variable sulfur cycling *Chloroflexus* were well represented (Blank et al., 2002; Chen et al., 2006; Eder and Huber, 2002; Huber et al., 1998; Stamps et al., 2014; Tang et al., 2011; Thiel et al., 2014). Potential phototrophs represented the most numerous putative metabolism in the top 15 most abundant taxa. Phototrophic genera *Chloracidobacterium, Geitlerinema* PCC 8501, *Chloroflexus*, GBChlB, *Roseiflexus*, and *Leptococcus* JA-3-3Ab (formerly classified as *Synechococcus* sp. JA-3-3Ab, currently *Leptococcus yellowstonii* (Walter et al., 2017)) were all highly represented along the SC outflow channel (Bhaya et al., 2007; Liu et al., 2012; Tang et al., 2021; Tank and Bryant, 2015b; Thiel et al., 2014). Overall*, Leptococcus* JA-3-3Ab was the second most abundant genus in the DNA fraction and the most abundant genus in the 16S rRNA (rel. % cDNA abundance) analysis followed by *Meiothermus* and *Geitlerinema* PCC 8501 (Supplemental Figure S6). *Tepidimonas* and *Rivularia* PCC 7116, a cyanobacterium, round out the top five most abundant taxa in the rRNA data.

The percent relative abundances of the rRNA (cDNA) and rRNA gene (DNA) sequences of the top eight taxa were compared to assess the activity versus presence of the microbial taxa (Supplemental Figure S7). Although *Meiothermus, Leptococcus* and *Geitlerinema* were abundant in both cDNA and DNA, *Meiothermus* was proportionally more abundant in DNA, while *Leptococcus* and *Geitlerinema* were more abundant in cDNA. In this analysis, the proportional representation was interpreted as microbial transcriptional activity versus microbial representation; therefore, *Leptococcus* and *Geitlerinema* were transcriptionally more active than they were represented in the microbial community, while *Meiothermus* was less. *Rivularia* PCC 7116 sequences, a putative phototroph, displayed a similar trend as *Leptococcus* and *Geitlerinema*. These results could be attributed to the fact that samples were collected during the day, inducing higher activity of phototrophs; however, both *Chloracidobacterium* and *Chloroflexus* were less active than represented.

### 3.5 Microbial Taxa Influencing Community Dynamics

To investigate the taxonomic shifts of microorganisms occurring spatially and temporally, differential abundance tests were conducted. Pairwise tests were conducted between sequential locations or dates for the 16S rRNA gene and rRNA sequences at the genus level to determine which genera were differentially abundant/active (Supplemental Figures S8-11). Thirty-three different genera were determined to be differentially abundant in the rRNA gene fraction temporally, while 34 genera in the rRNA fraction were differentially abundant. Fifty-seven and 48 unique genera were spatially differentially abundant in the rRNA gene and rRNA fractions respectively. Across all tests, 20 different genera were found to be differentially abundant in all instances. *Roseiflexus*, A4b family, *Thermus*, and *Lewinella* were the only taxa in the top 15 not to be differentially abundant in all four tests.

Beta diversity analysis revealed how variable and dynamic microbial communities at SC can be temporally and spatially. To better understand which taxa were driving these community shifts, a SIMPER analysis was conducted at the genus level (Clarke, 1993). In total, ASVs in 104 instances were determined to significantly contribute to dissimilarity between samples across time, belonging to thirty unique genera. No ASVs significantly contributed to spatial dissimilarity between samples. Of the top 15 most abundant taxa, only *Thermocrinis* and *Thermus* were not identified as significant in the analysis.

To determine the temporal and spatial dynamics of key microorganisms at SC, genera determined to be statistically significant in both the differential abundance and SIMPER analysis were subset and examined (Figure 6). These taxa were the most temporal-spatially dynamic and acted as microbial drivers for community shifts at SC. Fifteen genera were determined to be both significant drivers of community change and dynamic temporal-spatially. To better decipher the observed dynamics of these microorganisms, the most abundant ASV for each genus was run through NCBI BLAST where the most similar cultured representatives were researched to determine the probable and optimal putative metabolism associated with the ASV. This work revealed five general metabolisms in the 15 dynamic microorganisms: oxygenic and anoxygenic photosynthesis, aerobic anoxygenic photosynthesis, chemoautotrophy, and heterotrophy. *Leptococcus* JA-3-3Ab, *Geitlerinema* PCC-8501, *Leptolyngbya* FYG, and *Pseudanabaenaceae* family representatives predominantly grow as oxygenic phototrophs (OPs, Bhaya et al., 2007; Bohunická et al., 2011; Bosak et al., 2012a; Cordeiro et al., 2020; Momper et al., 2019; Tang et al., 2021), while *Chloroflexus* and *Chloracidobacterium* predominantly grow as anoxygenic phototrophs (APs, Tang et al., 2011; Tank and Bryant, 2015b, 2015a; Thiel et al., 2014). Aerobic anoxygenic phototrophs (AAP) were represented by *GBChlB* and *Tabrizicola* (Liu et al., 2012; Tarhriz et al., 2013, 2019). ASVs belonging to the *Methylacidiphilaceae* family, chemoautotrophs, shared 99.47% similarity to the methanotrophs *Methylacidiphilum infernorum* and *Methylacidiphilum kamchatkense* (Hou et al., 2008; Kruse et al., 2019). The remaining taxa *Raineya, Meiothermus, Tepidimonas, Thermoflexibacter, Saprospiraceae* Family, and WD2101 soil group family were all identified as heterotrophs (Albuquerque et al., 2009, 2018; Chen et al., 2006; Dedysh et al., 2021; Hahnke et al., 2016; Lewin and Lounsbery, 1969; McIlroy and Nielsen, 2014; Tindall et al., 2010; Xia et al., 2008).

**Figure 6.**
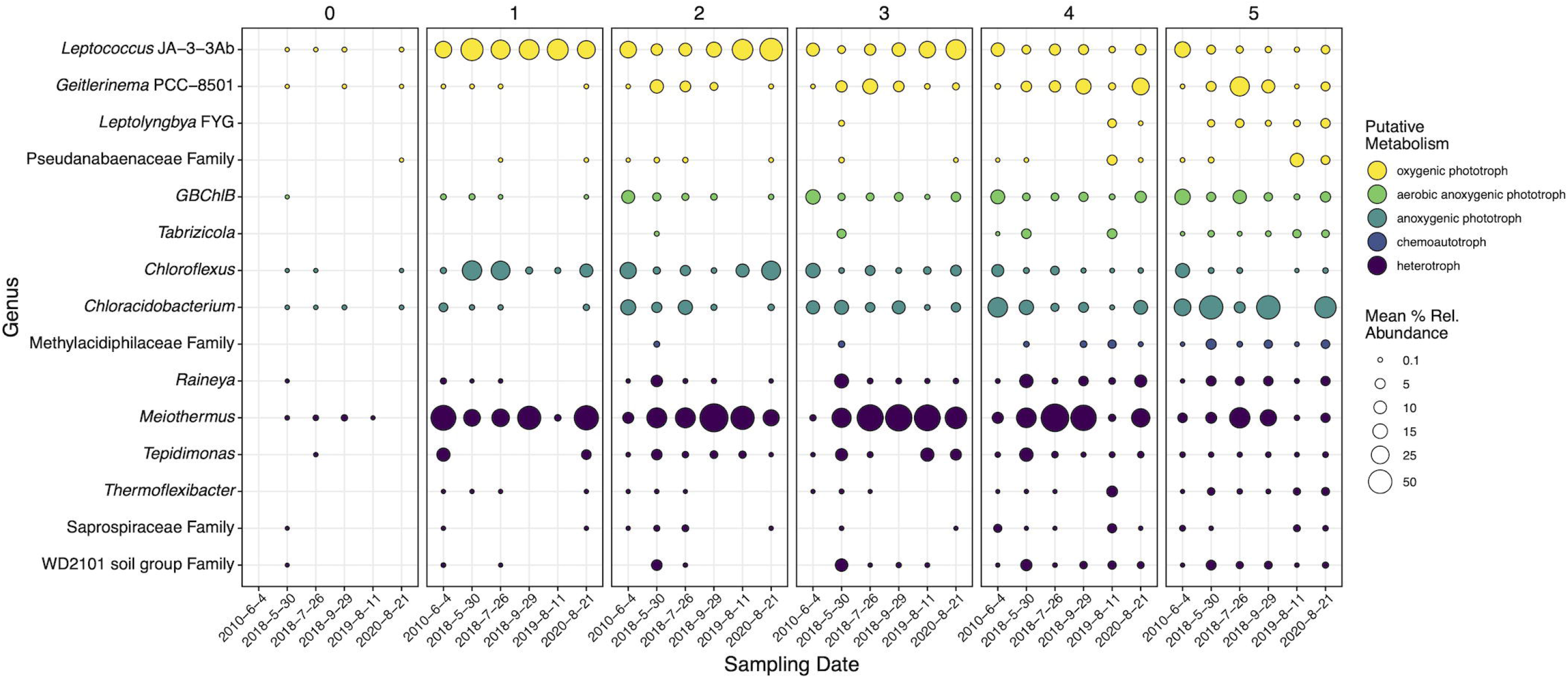
16S rRNA gene mean percent relative abundances of genera (or best classification) determined to be statistically significant by SIMPER, DNA differential abundance analysis, and cDNA differential abundance analysis. Panels are faceted by distance (m) from the hydrothermal source. Circle sizes indicate mean percent relative abundance. Colors indicate general putative metabolisms associated with the most abundant ASVs in each genus according to NCBI BLAST searches and literature review. Dates are shown as year-month-day.

Comparatively, multiple dynamics were displayed within each putative metabolism both temporally and spatially. Heterotrophs *Raineya*, *Thermoflexibacter*, *Saprospiraceae* family, and WD2101 soil group family increased in abundance down the outflow; however, *Meiothermus* and *Tepidimonas* became abundant (>50% or >10% respectively) at 1 m and persisted until 4 and 5 m, respectively. *Meiothermus* and *Tepidimonas* often coexisted at steady abundances temporally and spatially, with *Meiothermus* often ranging from 20 to 70% and *Tepidimonas* from 2 to 15%. *Raineya, Thermoflexibacter*, *Saprospiraceae* family, and WD2101 soil group family exhibited the same trend in the 16S rRNA sequencing data (Supplemental Figures S12), becoming more active with increasing distance from the hydrothermal source. *Meiothermus* was more abundant closer to the source (~50%) and steadily decreased to <10% down the transect in the rRNA fraction, a trend that was not present in the rRNA gene data. Also, *Meiothermus* steadily decreased in rRNA abundance temporally across all distances (except 1 m), while *Tepidimonas* mirrored its 16S rRNA gene trend.

Of the four dynamic OPs, all except *Leptococcus* JA-3-3Ab increased in abundance down the transect. *Geitlerinema* PCC-8501 and *Leptococcus* JA-3-3Ab were the two most abundant phototrophs, ~20% and ~40% respectively, and exhibited spatially inverse trends that overlapped between 2 and 5 m. *Leptolyngbya* FYG and *Pseudanabaenaceae* family displayed similar trends spatially; however, *Pseudanabaenaceae* was mostly present in 2019 at 4 and 5 m. AP microorganisms displayed inverse spatial abundances, *Chloracidobacterium* increased from <1% to ~50% abundance after 5 m, while *Chloroflexus* decreased from 30% at 1 m to <5% at 5 m. Unlike the OPs and APs, the two AAPs exhibit the same spatial trend, increasing with increasing distance from the hydrothermal source. *GBChlB* increased from <1% abundance to >30% while *Tabrizicola* increased from 0% to ~4%. *Tabrizicola* displayed pulses of increased abundance in May 2018 and 2019 leading to increased abundance closer to the source, while *GBChlB* decreased in overall abundance from 20% in 2010 to ~5% in 2020. Unlike the dynamic heterotrophic taxa, phototrophs displayed similar trends in both the rRNA and rRNA gene sequencing data.

## 4 Discussion

Actively silicifying hydrothermal systems are the modern analogs of ancient silicified rock deposits and the microfossils preserved therein. To determine the geomicrobiological dynamics of an actively silicifying hydrothermal system we tracked both microbial communities via 16S rRNA/rRNA gene sequencing of biofilms as well as aqueous geochemistry along the geyser outflow channel. Overall, geochemical conditions were steady at individual locations with minimal variation over a decade, but distinct microbial gradients likely tied to temperature existed as the waters moved from the hydrothermal source to the sampled transect terminus. Contrary to geochemical stability, microbial communities exhibited temporal dynamics resulting from fluctuations in the relative abundances of key microorganisms. Furthermore, microbial taxa exhibited genera-specific spatial stratification. Conceptually, the abiotic impactors that influence degassing, cooling, and geochemical oxidation remained constant due to the open nature of the system (Nordstrom et al., 2005). Yet, biotic influences such as biological sulfide or arsenic oxidation could vary as indicated by the dynamic nature of the microbial communities. Therefore, we postulate that temporal geochemical outflow stability is attributed to consistent abiotic forces and microbial community metabolic redundancy. Further investigation is needed to disentangle the extent and magnitude of these effects on outflow waters. By conducting a paired temporal and spatial study, we show that the limits of geological end-point studies may be constrained to silicified microfossils and their proximity to a hydrothermal vent source.

In hot springs, differences in geochemistry (e.g., aqueous chemistry, temperature, dissolved oxygen, etc.) play a large role in dictating the possible microbial communities present. Any shifts in geochemistry, particularly temperature, over time could alter the observable microbial communities. The geochemistry was largely stable spatially at SC, which is indicative of the complex interplay between abiotic and potential biotic forces with similar trends observed at other hot spring systems (Figures 2 and S4, Costa et al., 2009; Swingley et al., 2012). The oligotrophic source waters at SC remained steady throughout the ten-year sampling period, showing minimal significant changes in the major analytes (Table 1). The stability of the source is largely attributable to a predominantly hydrothermal hydrological reservoir feeding SC (Nordstrom et al., 2009). Waters that reside deeper in the subsurface receive minimal meteoric water input and therefore have less influence from heterogeneous shallow subsurface geology and precipitation variability (Fournier, 1989). As the hydrothermal waters cool along the SC outflow from 0 to 1 m, a major temperature threshold was crossed (72/73°C, the upper limit of photosynthesis, (Brock, 1978; Kees et al., 2022)) allowing for the proliferation of phototrophs such as *Leptococcus* and *Geitlerinema*. Such a major shift in detectable microbial communities indicates that temperature is critical in the establishment of fundamental niches at SC, a precedent previously established at other biomat locations (Bennett et al., 2020; Wang et al., 2013). Temperature also appears important for the shift in major members of the community as a pulse in increased abundance of mesophilic *Meiothermus, Tepidimonas, Raineya*, and the WD2102 soil group in May 2018 corresponds with cooler temperatures, especially at 2, 3, and 4 m (Figures 2 and 5). *Meiothermus* and *Tepidimonas taiwanensis* possess optimal growth temperatures of 60 and 55°C respectively (Albuquerque et al., 2009; Chen et al., 2006; Tindall et al., 2010), and are most abundant between 1 and 5 m for *Meiothermus* or 1 and 4 m for *Tepidimonas* (Figures 5 and 6). Our results indicate a narrower temperature range for *Tepidimonas* growth than previous laboratory experiments with abundance plummeting below 40°C (Chen et al., 2006). Furthering thermally induced spatial stratification, *Raineya* (T_opt_ = 50°C) and WD2101 soil group (observed range = 47-50°C) taxa became more abundant down transect beginning at 2 and 3 m respectively (Albuquerque et al., 2018; Dedysh et al., 2021).

While temperature is critical in the establishment of niches within SC, nutrient limitation likely controls maximum productivity (i.e., biofilm thickness) down channel and may also influence the presence of microorganisms adapted to nutrient limiting conditions. Measured phosphorus and nitrogen species remained consistently low or below our quantification limits along the outflow, suggesting that SC is limited in both phosphorus and nitrogen. While nitrogen could be sourced from atmospheric nitrogen and N2 fixing phototrophs, it remains unclear how phosphorus is sourced to sustain a thriving microbial biofilm. There is precedence for microbial growth in oligotrophic environments under highly limiting phosphate conditions (Erb et al., 2012). Steep Cone is geographically isolated as an elevated platform in the middle of Sentinel Meadow and hydrothermally sourced high-chloride springs have been shown to receive minimal subsurface phosphorus (Stauffer and Thompson, 1978). Dust and animal inputs (e.g. insects, bison) likely represent the primary inputs of phosphorus into the system as has been shown previously in other phosphorus limited spring systems (Strumness, 2006). Inorganic carbon (DIC) concentrations remained consistently high down the transect, while organic carbon (DOC) concentrations remained consistently low. This is surprising considering the carbon requirement for actively growing phototrophic community members, which due to low initial DOC input would predominantly grow photoautotrophically. Evidence of robust phototrophic microorganisms would inherently introduce more organic carbon into the system down transect, which is not observed. We surmise that phototrophically fixed carbon remains bound in the biofilms and that heterotrophs consume this organic carbon prior to dissolution into the overlying waters. While the steady concentration of DIC along the outflow would indicate minimal autotrophic consumption, chloride concentrations down transect indicate evaporation trends (Swingley et al., 2012), which would in turn continuously act as a concentration mechanism offsetting any metabolic consumption of DIC. The requirement for carbon and nitrogen fixation and minimal input of bioavailable phosphorus likely limit productivity at SC.

With a limited pool of nutrients, the biofilms at SC are thin and adhere to the mineral crust at the surface. The sampled biofilm at SC was 1 to 3 mm thick, much thinner than the classically studied stratified biomats such as Mushroom Spring (YNP), which are multiple cm thick (Thiel et al., 2017; Wörmer et al., 2020). Despite how thin the sampled biomat was, microbial abundance data and putative metabolisms points towards a depth stratified biofilm (Figure 6), especially further down the transect. In accordance with previous studies of stratified biofilms and biomats, OPs and APs occupy the top and bottom layers of the mat respectively (Boomer et al., 2009; Thiel et al., 2017; Wörmer et al., 2020). We propose AAPs occupy the interface between OPs and APs in the biofilm at SC, and potentially fluctuate higher or lower in the biofilm based upon metabolic needs. AAPs have been shown to be aerobic, possess bacteriochlorophyll *a*, and the versatility to thrive in oligotrophic environments (Kolber et al., 2001; Suyama et al., 2002). *Tabrizicola aquatica*, an AAP, was shown to be metabolically capable of chemotrophic and phototrophic growth via bacteriochlorophyll *a*, yet it requires oxygen and cannot grow under anaerobic phototrophic conditions (Tarhriz et al., 2019), which would spatially place *Tabrizicola* found at SC between the OPs and APs based upon growth requirements.

With the fundamental niches of SC established by temperature gradients and nutrient limitation we also observed microorganisms that were putatively linked to three general metabolic categories: phototrophy, heterotrophy, and sulfur cycling microorganisms. Due to the oligotrophic source waters, phototrophs such as *Leptococcus*, *Geitlerinema*, and *Chloroflexus* act as the primary producers in the system after 1 m, while *Thermocrinis* and *Thermus*, the only inhabitants at the source, are likely thriving via lithotrophic metabolisms under the extreme temperatures, low DOC/high DIC waters at the source (Caldwell et al., 2010; Zhou et al., 2020). Additionally, the presence of heterotrophs corresponds to locations exhibiting the proliferation of phototrophs, linking heterotroph reliance on phototrophs as the primary producers of reduced carbon substrates at SC. Heterotroph and phototroph abundances combined with DIC/DOC data further support the above hypothesis regarding carbon consumption (Figures 2 and 6).

The dynamics between phototrophic microorganisms at SC are more complex than the heterotrophic component. The temporal trends and drivers of phototrophic shifts are separated by the potential type of photosynthesis carried out by different microorganisms that drive community shifts (Figure 6). For example, the cyanobacteria *Leptococcus* and *Geilterinema* display an inverse spatial distribution, and both are present in high abundances across time from 2 to 4 m, indicating potential niche competition. Yet, *Leptococcus* genera have been shown to be capable of photoautotrophy, photoheterotrophy, and chemoorganotrophy (Bhaya et al., 2007), while *Geitlerinema* have the potential to perform both oxygenic photosynthesis or anoxygenic photosynthesis using H2S as an electron donor (Hamilton et al., 2018; Momper et al., 2019; Tang et al., 2021). *Leptolynbya* mimic the metabolic capabilities of *Geitlerinema* (Bosak et al., 2012b; Hamilton et al., 2018). Similarly, AAPs display location and temporal overlap at SC. AAPs have been shown to be metabolically diverse and highly adaptive, capable of compensating for low DOC concentrations by increasing bacteriochlorophyll *a* and photosynthetic production (Beatty et al., 2002; Suyama et al., 2002; Tarhriz et al., 2019). APs *Chloroflexus* and *Chloracidobacterium* display temporal and spatial overlap in the middle reaches of the transect (2-4 m) and earlier sampling dates (Figure 6, 2010 and 2018s). While *Chloracidobacterium* display minimal metabolic flexibility, primarily growing only anoxygenic photoheterotrophically (Tank and Bryant, 2015b, 2015a), *Chloroflexus* have been shown to be metabolically robust, capable of anoxygenic photoautotrophy, photoheterotrophy, aerobic chemoorganotrophy, and capable of utilizing various sulfur species (Nübel et al., 2001; Tang et al., 2011; Thiel et al., 2014). Therefore, metabolic flexibility may allow the identified OPs, AAPs, and APs to thrive at the same locations and times at SC. This flexibility does appear to have its limits as we observed ebb and flow of the 15 genera responsible for community shifts over the sampling period.

Upon inspection of the distribution and abundances of various phototrophs at SC, both growth down transect and biofilm stratification relationships emerge (Figure 6). In agreement with heterotrophic spatial stratification, phototrophs expectedly distribute along their optimal growth temperature ranges. For example, *Chloroflexus* and *Chloracidobacterium*, both potential anoxygenic phototrophs (AP), were shown to have optimal growth ranges of 52-60 and 44-58°C respectively (Pierson and Castenholz, 1974; Tank and Bryant, 2015b), and abundance data illustrates an inverse relationship with *Chloroflexus* decreasing (>30% to <10%) while *Chloracidobacterium* increases (<5% to ~ 50%) from 1 to 5 m (Figure 5). The spatial abundance patterns of the oxygenic phototrophic (OP) cyanobacteria *Leptococcus* and *Geitlerinema* mirror the APs inverse relationship with their closest representatives via BLAST exhibiting optimal growth temperature of 62 and 40°C respectively (Allewalt et al., 2006; Tang et al., 2021). In all dynamic genera, taxa shown to have higher growth temperatures were found to have greater abundance than similar taxa with lower growth temperatures, despite the measured outflow temperatures being complementary to the latter’s optimum. These results further support that spatial growth is being driven by the temperature gradient along the outflow at SC.

While no microbial taxa were determined to be significant drivers spatially, numerous genera shifted community diversity temporally. Here, we contend that a combination of niche competition, potential metabolic flexibility, and geochemistry contributes to the proliferation and fall of significant taxa that drive community change. *Meiothermus*, *Tepidimonas*, and *Raineya* were shown to occupy similar locations spatially, possess similar putative metabolisms, but often coexist at the same times in abundances greater than 10% (Figures 5 and 6, 2010, May 2018, 2019, 2020). Their similar lifestyles suggest niche competition in the oligotrophic waters at SC. *Meiothermus* and *Raineya* species have been shown to be aerobic chemoorganotrophs, while *Tepidimonas* are aerobic chemolithoheterotrophs capable of oxidizing reduced sulfur compounds to sulfate (Albuquerque et al., 2018; Moreira et al., 2000; Tindall et al., 2010). We observed sulfur oxidation along the outflow to predominately SO_4_^2-^ after 5 m (Figure 2). This potential metabolic flexibility may allow *Tepidimonas* to thrive concurrently with other abundant mesophilic heterotrophs While previous work at a similar hot spring system attributed a majority of sulfur oxidation to abiotic forces (Nordstrom et al., 2005), sulfur oxidizing microorganisms are present along the outflow transect and likely contribute to the observed sulfate concentrations.

One final, possibly cryptic metabolic niche at SC is arsenic cycling. Arsenic concentration increased down the transect; the driver for the observed increase is supported by evaporative concentration not arsenic dissolution, which has been shown not to be present at the SC deposit (Gangidine et al., 2020; Yokoyama et al., 2004). Of the taxa present at SC, only *Thermocrinis* and *Thermus* have been linked to metabolic arsenic cycling (Caldwell et al., 2010; Hartig et al., 2014; Zhou et al., 2020) and were observed at the hydrothermal vent but not further down the transect. The lack of metabolically active nutrients within the SC source waters and comparatively high arsenic concentrations indicates a potential unexplored (and undetected) metabolic niche for arsenic cycling microorganisms at SC, especially anoxygenic photosynthesis coupled to arsenic oxidation (Kulp et al., 2008; Oremland and Stolz, 2003).

While specific microbial taxa are important at SC, the richness and distribution of microbial communities present throughout the geyser system over time are critical to understanding how the system evolves. Richness analysis illustrated the proliferation of microbial taxa *(i.e*., increased number of observed taxa) down the outflow, indicating the importance of cooling waters in the amount of diversity at any given location (Figure 3). In essence, as the hydrothermal waters cool, less selective pressure is placed upon the microbial communities allowing for a greater number of unique taxa to find a foothold and live. Similarly, evenness increased from source to terminus plateauing at 2 m and ~50°C (Figure 3). These results indicate that the ceiling on evenness is the selective temperature gradient and oligotrophic system acting upon the microbial communities. The oligotrophic water of the spring indicates that while more unique taxa can live as the temperature regime cools, they are competing for the same limited resources *(e.g.*, organic carbon, nitrogen, and phosphorus), resulting in only a few metabolically viable niches for new taxa. Sulfate concentrations similarly stagnate after 2 m indicating an oxidized environment (Figure 2, Nordstrom et al., 2005), causing marginalization of sulfate oxidative metabolisms. Therefore, after 2 m and limiting resources, there is a dominant taxon that occupies any given niche (i.e., *Leptococcus* or *Meiothermus* for 2 - 4 m, Figure 6) that is the most fit given the environmental conditions, while other less fit taxa and less abundant taxa compete over remaining nutrients, a dynamic that leads to increased richness and reduced evenness. We posit that after 2 m at SC, there are four generalized niches for most microorganisms to thrive: OP, AAP, AP, and heterotrophy.

Increasing richness and the observed plateau of evenness at SC elucidated the unique interplay between selective pressures and limitations in oligotrophic environments, while beta diversity metrics provide similar insights between entire microbial communities (Figure 4). The microbial communities at SC not only became more taxonomically rich along the exit, but also became less similar (Figure 4B). As microbial communities increased in distance from one another, geochemical parameters and temperature were likewise drastically different *(e.g.*, significantly cooler water temperatures from source to terminus). It is therefore no surprise that the microbial communities at the boiling hydrothermal source were 100% different via Bray-Curtis comparison from the microbial communities 5 m away from the source and ~55°C cooler. At intermediate distances from the source, Bray-Curtis dissimilarity also decreased (communities became more similar) with increasing distance between samples from 1 to 5 m. We attribute this to the presence of many taxa shared across the intermediate sample distances, while the end-member distances illustrate the extremes of the sampled thermal gradient. No ASVs were determined to significantly shift the microbial communities along the outflow channel; thus, the distinct decrease in similarity along the outflow further supports temperature driven spatial stratification of microbial communities. Analyzing beta diversity variance trends at SC also exhibited how taking only single timepoint *(i.e.*, endpoint) measurements may miss the sporadic observed temporal dynamics of a hot spring (Figure 4A). Variance across all locations increased as the number of days after initial sampling increased, illustrating diverging similarities locationally. Specifically, SIMPER analysis for 2019 and 2020 sampling events implicate abundance shifts in *Meiothermus, Geitlerinema* PCC-8501, and *Tuwongella* (a potential heterotroph, Seeger et al., 2017) as the main contributors to community change. *Meiothermus* was highly abundant across most locations in September 2018 and 2020 samples, but only present at 2 and 3 m in 2019. *Geitlerinema* similarly disappears and reappears over the same period. Therefore, significant changes in temporal microbial diversity are driven by abundance shifts of the top 15 most abundant taxa over yearly time scales, potentially influencing the findings of studies that may only come upon a silicifying hydrothermal system after it has lithified.

Further research regarding silicification of microorganisms is ongoing; however, the dynamic nature of the microbial communities and dominant taxa comprising these communities is undeniable and should give pause to any research attempting to draw firm microbiological conclusions from entombed microorganisms. Microbial communities differentiate along outflow channels and temporally, causing end point conclusions of the silicified rock record to become constrained. One common method to identify lithified or ancient microbial communities is by scanning electron microscopy (SEM) and making inferences based on cell type *(e.g.*, filaments, rods, etc.) In agreement with previous work that identified filaments as the predominant morphology in silicified environments, our taxonomic and SEM work shows similar trends (Figures S2 and S3). Genera such as *Geitlerinema*, *Leptolyngbya*, and *Leptococcus* are highly abundant and potentially filamentous (Bhaya et al., 2007; Bohunická et al., 2011; Hamilton et al., 2018). Similarly, the biofilms observed with SEM imaging were dominated by filamentous structures amidst a thick coat of EPS. Taxa associated with rod-shaped morphologies are also present and abundant such as *Meiothermus*, *Thermocrinis*, and *Tabrizicola*. These taxa have also been shown to form filaments under certain conditions (Albuquerque et al., 2009; Eder and Huber, 2002; Nobre et al., 1996; Tarhriz et al., 2019) potentially confounding any results that may only draw from SEM to establish ancient microbial community composition. Fluid flow induced filament/trichome formation by rod or coccoid cells along the outflow channel may also explain the relative dearth of nonfilamentous morphologies visible in the biofilm or silicified microfossils, a phenomenon previously documented in *Staphylococcus aureus* (Kevin Kim et al., 2014). 16S rRNA and rRNA gene analysis indicate that the mere presence of microbial taxa may not be indicative of their impact on the ecosystem (Figures S7 and S12). For example, *Meiothermus* is more abundant in the rRNA gene fraction than the rRNA, while *Leptococcus* is more abundant in the rRNA fraction, indicating *Leptococcus* is disproportionately more active in the system than it is represented. We show that while geochemical stability exists over large periods of time, the microbial communities in silicifying hot spring systems can be highly dynamic over yearly periods. As a result, we emphasize that caution should be taken when extrapolating end-point findings of the silicified rock record to predict ancient ecosystems.

## 5 Conclusion

This study employed 16S rRNA and rRNA gene sequencing in combination with geochemical analysis to characterize both the temporal and spatial dynamics of microbial communities along the outflow of an actively silicifying hydrothermal system. We determined the distinct drivers of microbial community change temporally and spatially, especially how microorganisms arise and fall through both time and space. Our results indicate that community stratification is primarily due to temperature gradients along the outflow channel. Hyperthermophiles such as *Thermocrinis* dominate the source biofilm communities while thermotolerant microorganisms proliferate with *Leptococcus* and *Meiothermus* being major constituents at the middle outflow locations and finally giving way to mesophilic taxa at the end of the sampled transect. Phototrophy likely contributes the majority of observed organic carbon and places fundamental limits upon the total possible productivity of the biofilms. The results further indicate that the microbial communities undergo large changes yearly, driven by the ebb and flow of dominant taxa along the outflow. Geochemical and taxonomic analysis indicate that limited metabolic niches are present along the oligotrophic outflow. Furthermore, distinct phototrophic community members such as *Leptococcus*, *Tabrizicola*, and *Chloracidobacterium* are dominant representatives of distinct phototrophs acting as the primary producers for the outflow system which support growth of thermotolerant heterotrophs such as *Tepidimonas*. Critically we caution that the dynamic nature of microbial communities within a hydrothermal ecosystem could influence our ability to interpret the silicified rock record.

## 6 Data availability statement

The datasets and R code for data processing may be found in online repositories. Sequencing raw read are available through NCBI sequencing read archive, accession number PRJNA931285.

## 7 Author Contributions

KR led the study design and focus. SU conducted field sampling in 2010, while KR led all subsequent field campaigns with aid from BS and JS. Laboratory work and results processing was conducted by KR. KR analysed results. GV aided in data processing. KR wrote the manuscript and all authors reviewed, edited, and approved the submitted article.

## 8 Funding

This research was funded by a grant from The Edna Bailey Sussman Fund and The Geological Society of America awarded to KR. Additional funds were provided by a NASA Exobiology grant (80NSSC19K0479) awarded to JS.

## 9 Acknowledgements

We would like to thank Annie Carlson at the Yellowstone Center for Resources for help and guidance regarding sampling and permitting for Yellowstone National Park. All samples were collected under Yellowstone permit number YELL-5664 to JS. Field work was supported by Eilish Spear, Alex Honeyman, Michael Vega, Kevin Chan, Chris Trivedi, Aiyana Spear, and Emma Harrison.

## 10 Conflict of interest

The authors declare that the research was conducted in the absence of any commercial or financial relationships that could be construed as a potential conflict of interest.

## 11 Supplementary material

The Supplementary Material for this article can be found online at: (link)

## 12 Abbreviations

SC: Steep Cone Geyser
DIC: dissolved inorganic carbon
DOC: dissolved organic carbon
ASV: amplicon sequence variant
EPS: extracellular polymeric substances
LOQ: limit of quantification
SIMPER: similarity percentages
OP: oxygenic phototroph
AP: anoxygenic phototroph
AAP: aerobic anoxygenic phototroph
SEM: scanning electron microscope

## Notes

### Competing Interest Statement

The authors have declared no competing interest.

