## Supplemental Material for "Spatial and Temporal Dynamics at an Actively Silicifying Hydrothermal System"

### *Supplementary Material*

#### 1 Supplementary Tables

**Supplementary Table 1.** Sampling summary for major types of analysis/quantification for each sampling time point. Dates are shown as year-month-day.

| Sampling Date | 2010-6-4 | 2017-8-18 | 2018-5-30 | 2018-7-26 | 2018-9-29 | 2019-8-9 | 2020-8-21 |
|---|---|---|---|---|---|---|---|
| **16S rRNA Gene** | x | x | x | x | x | x | x |
| **16S rRNA** | - | - | - | - | x | x | x |
| **IC** | - | x | x | x | x | x | x |
| **ICP** | - | x | x | x | x | x | x |
| **pH** | - | x | x | x | x | x | x |
| **Temperature** | - | x | x | x | x | x | x |
| **DIC/DOC** | - | - | - | x | x | x | x |
| **Notes** | No samples at 0m | Only sampled spring source | | No DIC/DOC sample at 3m | | DIC/DOC vials at 4 and 5m broke in transit | |

**Supplementary Table 2.** Alpha diversity pairwise statistical tests results comparing group1 and group2 entries for each row, either by date or distance from spring source in meters. The Kruskal-Wallis and pairwise Wilcoxon tests were used to determine alpha diversity statistical significances (adjusted *p*-value < 0.05), along with the Wilcoxon effect size. Only statistically significant results are shown (adjusted *p* < 0.05). * denotes *p* < 0.05, ** denotes *p* < 0.005, *** denotes *p* < 0.0005, **** denotes *p* < 0.00005. Mag. denotes magnitude of effect size and stat denotes statistic

| metric | category | group 1 | group 2 | n 1 | n 2 | stat | p | p.adj | p.adj. signif | effect size | mag. |
|---|---|---|---|---|---|---|---|---|---|---|---|
| **evenness** | date | 4-Jun-10 | 18-Aug-17 | 14 | 2 | 28 | 0.017 | 0.044 | * | 0.556 | large |
| **evenness** | date | 4-Jun-10 | 30-May-18 | 14 | 16 | 185 | 0.002 | 0.01 | * | 0.554 | large |
| **evenness** | date | 4-Jun-10 | 21-Aug-20 | 14 | 16 | 196 | 2.3E-04 | 0.005 | ** | 0.638 | large |
| **evenness** | date | 18-Aug-17 | 26-Jul-18 | 2 | 16 | 0 | 0.013 | 0.039 | * | 0.53 | large |
| **evenness** | date | 18-Aug-17 | 29-Sep-18 | 2 | 16 | 0 | 0.013 | 0.039 | * | 0.53 | large |
| **evenness** | date | 30-May-18 | 29-Sep-18 | 16 | 16 | 48 | 0.002 | 0.01 | * | 0.533 | large |
| **evenness** | date | 26-Jul-18 | 21-Aug-20 | 16 | 16 | 195 | 0.011 | 0.039 | * | 0.446 | moderate |
| **evenness** | date | 29-Sep-18 | 21-Aug-20 | 16 | 16 | 215 | 0.001 | 0.007 | ** | 0.58 | large |
| **evenness** | distance | 0 meters | 4 meters | 7 | 18 | 14 | 0.002 | 0.019 | * | 0.593 | large |
| **evenness** | distance | 0 meters | 5 meters | 7 | 18 | 17 | 0.004 | 0.019 | * | 0.557 | large |
| **evenness** | distance | 0 meters | 1 meters | 7 | 17 | 23 | 0.02 | 0.042 | * | 0.473 | moderate |
| **evenness** | distance | 0 meters | 2 meters | 7 | 18 | 16 | 0.003 | 0.019 | * | 0.569 | large |
| **evenness** | distance | 0 meters | 3 meters | 7 | 18 | 19 | 0.006 | 0.023 | * | 0.533 | large |
| **evenness** | distance | 1 meters | 4 meters | 17 | 18 | 77 | 0.011 | 0.034 | * | 0.424 | moderate |
| **evenness** | distance | 1 meters | 5 meters | 17 | 18 | 81 | 0.017 | 0.042 | * | 0.402 | moderate |
| **richness** | date | 4-Jun-10 | 29-Sep-18 | 14 | 16 | 223 | 2.8E-08 | 5.78E-07 | **** | 0.842 | large |
| **richness** | date | 30-May-18 | 29-Sep-18 | 16 | 16 | 246 | 4.6E-07 | 3.24E-06 | **** | 0.786 | large |
| **richness** | date | 26-Jul-18 | 29-Sep-18 | 16 | 16 | 237 | 6.9E-06 | 3.62E-05 | **** | 0.726 | large |
| **richness** | date | 26-Jul-18 | 21-Aug-20 | 16 | 16 | 47 | 0.002 | 0.006 | ** | 0.54 | large |
| **richness** | date | 29-Sep-18 | 11-Aug-19 | 16 | 16 | 44 | 0.001 | 0.004 | ** | 0.56 | large |
| **richness** | date | 29-Sep-18 | 21-Aug-20 | 16 | 16 | 9 | 3.2E-07 | 3.24E-06 | **** | 0.793 | large |
| **richness** | distance | 0 meters | 4 meters | 7 | 18 | 21 | 0.009 | 0.035 | * | 0.508 | large |
| **richness** | distance | 0 meters | 5 meters | 7 | 18 | 14 | 0.002 | 0.009 | ** | 0.593 | large |
| **richness** | distance | 1 meters | 4 meters | 17 | 18 | 56 | 0.001 | 0.007 | ** | 0.541 | large |
| **richness** | distance | 1 meters | 5 meters | 17 | 18 | 48 | 3E-04 | 0.004 | ** | 0.586 | large |

**Supplementary Table 3.** Beta diversity pairwise statistical test results. Permutational analysis of variance (PERMANOVA) and pairwise comparisons (adjusted *p*-value < 0.05) were calculated using Vegan to test for statistical significance between beta diversity groupings across dates and locations (10000 permutations). Only statistically significant results are shown (adjusted $p < 0.05$). Pairwise comparisons were performed between entries in variable 1 column and variable 2 column.

| test. variable | p.value | adj.p.value | variable 1 | variable 2 | sig | R2 |
|---|---|---|---|---|---|---|
| **Distance from spring** | 0.019 | 0.019 | 4 meters | 3 meters | TRUE | 0.066 |
| **Distance from spring** | 0.032 | 0.032 | 2 meters | 3 meters | TRUE | 0.065 |
| **Sampling date** | 1.00E-04 | 1.00E-04 | 4-Jun-10 | 30-May-18 | TRUE | 0.234 |
| **Sampling date** | 1.00E-04 | 1.00E-04 | 4-Jun-10 | 26-Jul-18 | TRUE | 0.277 |
| **Sampling date** | 1.00E-04 | 1.00E-04 | 4-Jun-10 | 29-Sep-18 | TRUE | 0.184 |
| **Sampling date** | 1.00E-04 | 1.00E-04 | 4-Jun-10 | 11-Aug-19 | TRUE | 0.183 |
| **Sampling date** | 3.00E-04 | 3.00E-04 | 4-Jun-10 | 21-Aug-20 | TRUE | 0.14 |
| **Sampling date** | 0.042 | 0.042 | 30-May-18 | 26-Jul-18 | TRUE | 0.073 |
| **Sampling date** | 2.00E-04 | 2.00E-04 | 30-May-18 | 11-Aug-19 | TRUE | 0.153 |
| **Sampling date** | 2.00E-04 | 2.00E-04 | 30-May-18 | 21-Aug-20 | TRUE | 0.134 |
| **Sampling date** | 3.00E-04 | 3.00E-04 | 26-Jul-18 | 11-Aug-19 | TRUE | 0.137 |
| **Sampling date** | 1.00E-04 | 1.00E-04 | 26-Jul-18 | 21-Aug-20 | TRUE | 0.197 |
| **Sampling date** | 0.011 | 0.011 | 29-Sep-18 | 21-Aug-20 | TRUE | 0.099 |
| **Sampling date** | 0.004 | 0.004 | 11-Aug-19 | 21-Aug-20 | TRUE | 0.125 |

#### 2 Supplementary Figures

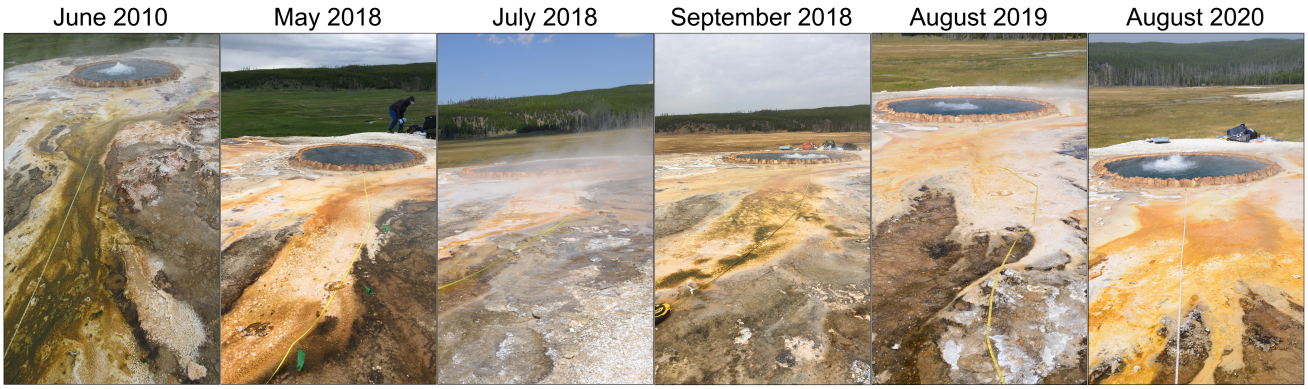


**Supplementary Figure 1.** Sampled outflow path at Steep Cone Geyser for each sampling date.

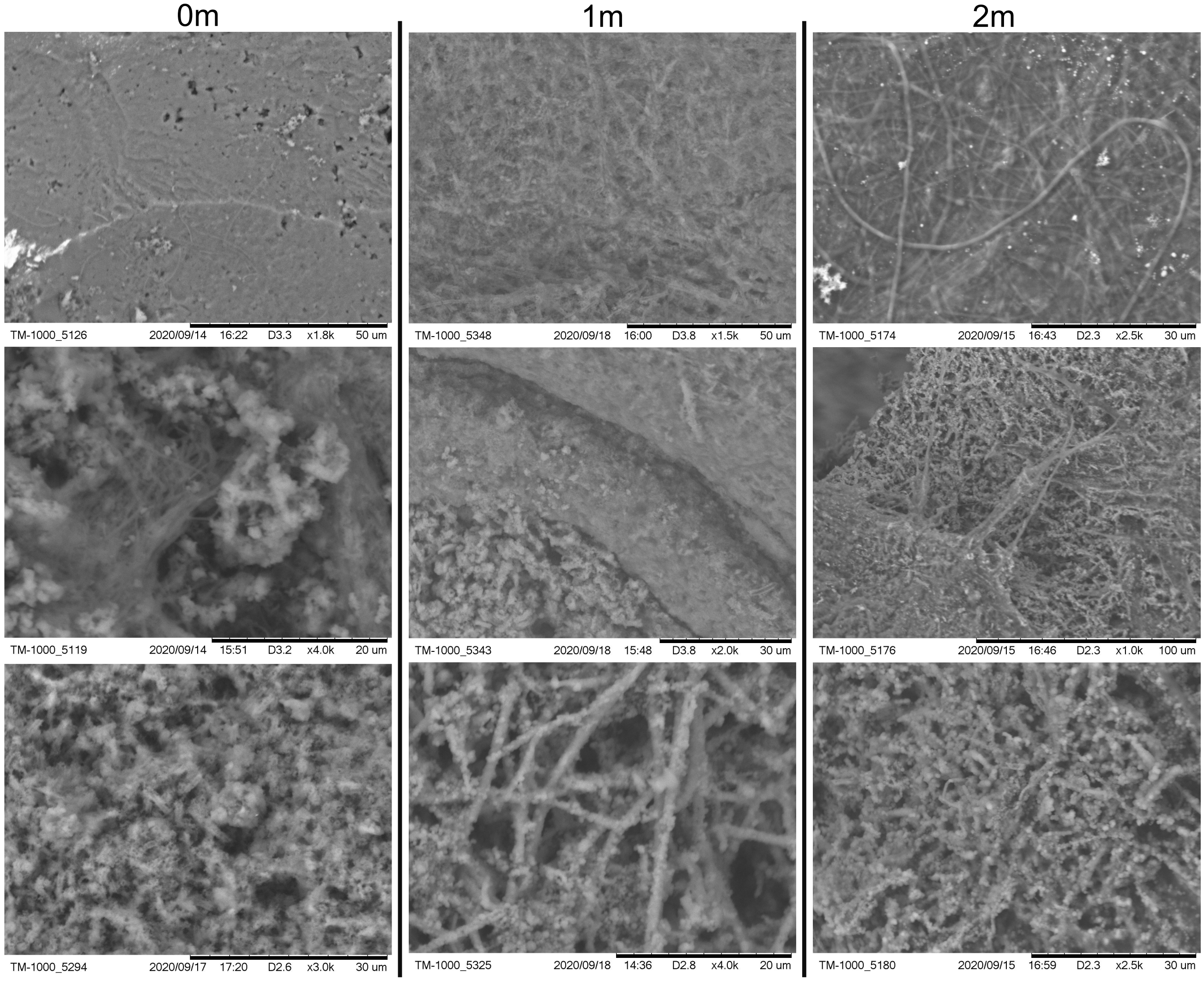


**Supplementary Figure 2.** Scanning electron microscope images taken of the Steep Cone Geyser outflow channel biofilm and underlying silicified matrix at 0, 1, and 2 meters. Top numbers indicate distance from the hydrothermal source the samples were taken at in meters. Each column corresponds to the listed distance. Images illustrate thin EPS and filament rich biofilm atop a fully silicified microbial matrix.

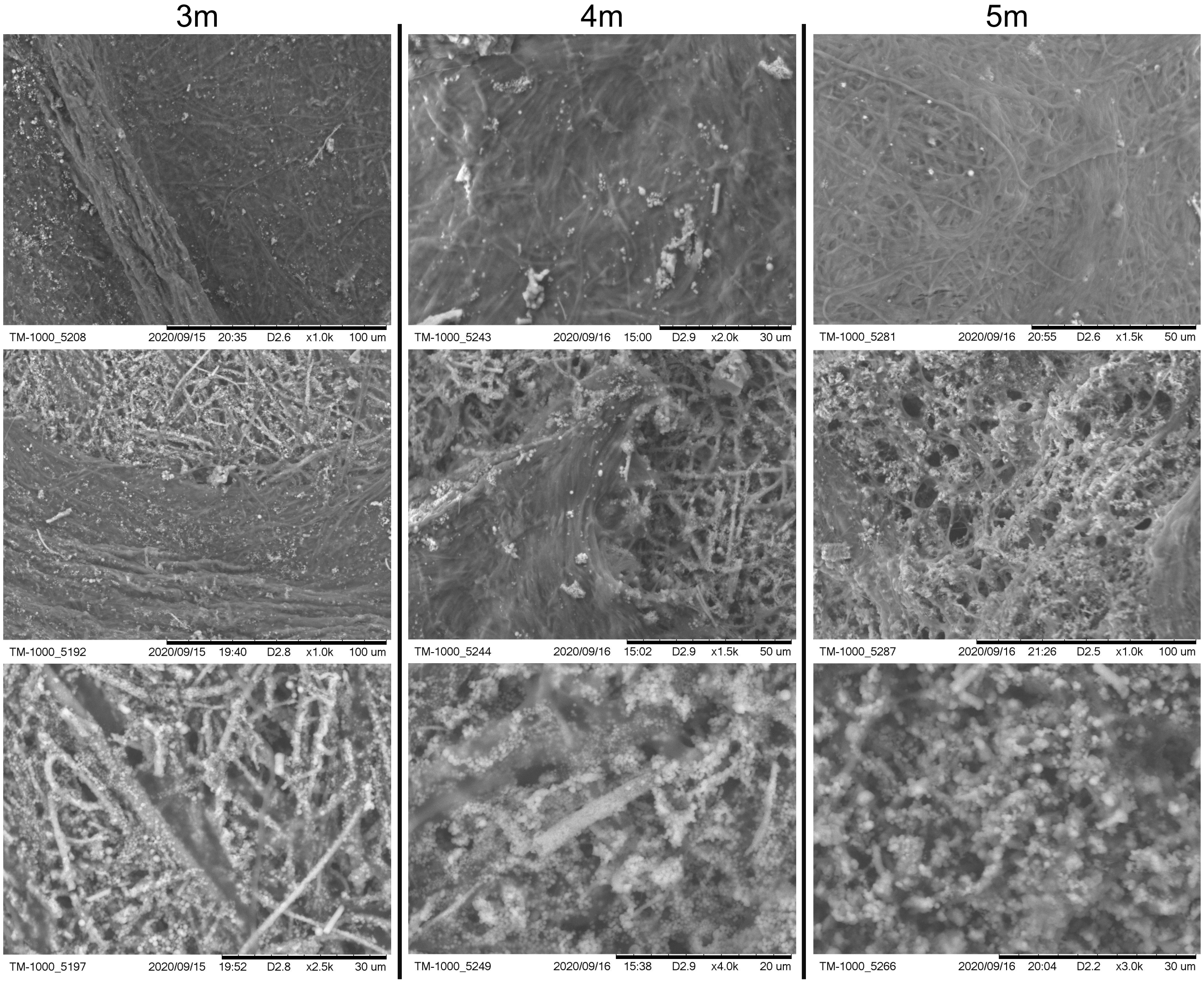


**Supplementary Figure 3.** Scanning electron microscope images taken of the Steep Cone Geyser outflow channel biofilm and underlying silicified matrix at 3, 4, and 5 meters. Top numbers indicate distance from the hydrothermal source the samples were taken at in meters. Each column corresponds to the listed distance. Images illustrate thin EPS and filament rich biofilm atop a fully silicified microbial matrix.

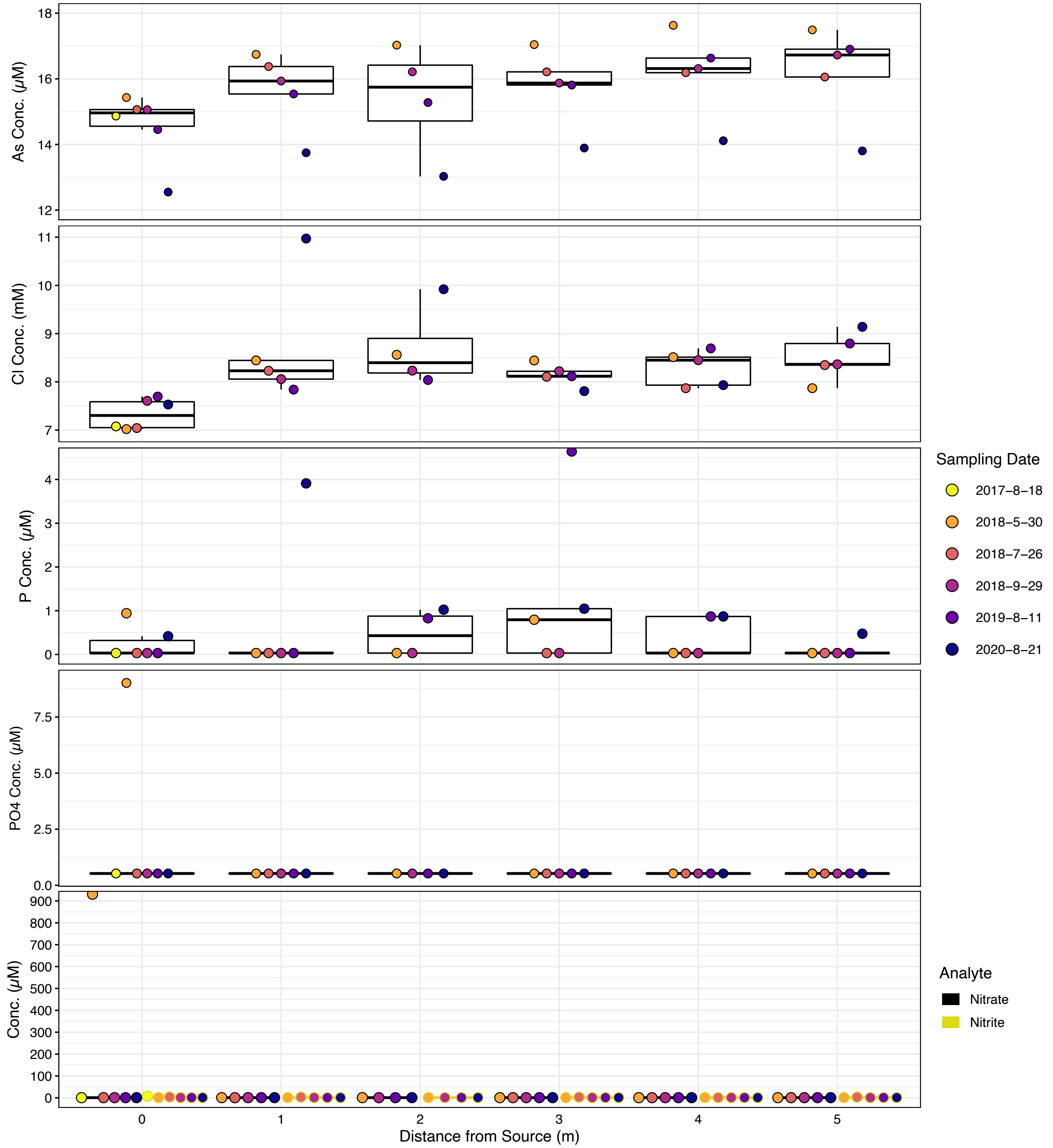


**Supplementary Figure 4.** Dot and box plot illustrating geochemical stability down the sampled transect across sampling dates of geochemical parameters of interest. Filled dot colors indicate sampling date while box colors indicate measured parameter. Dots are arranged from left to right, oldest to most recent samples respectively. Dates are shown as year-month-day.

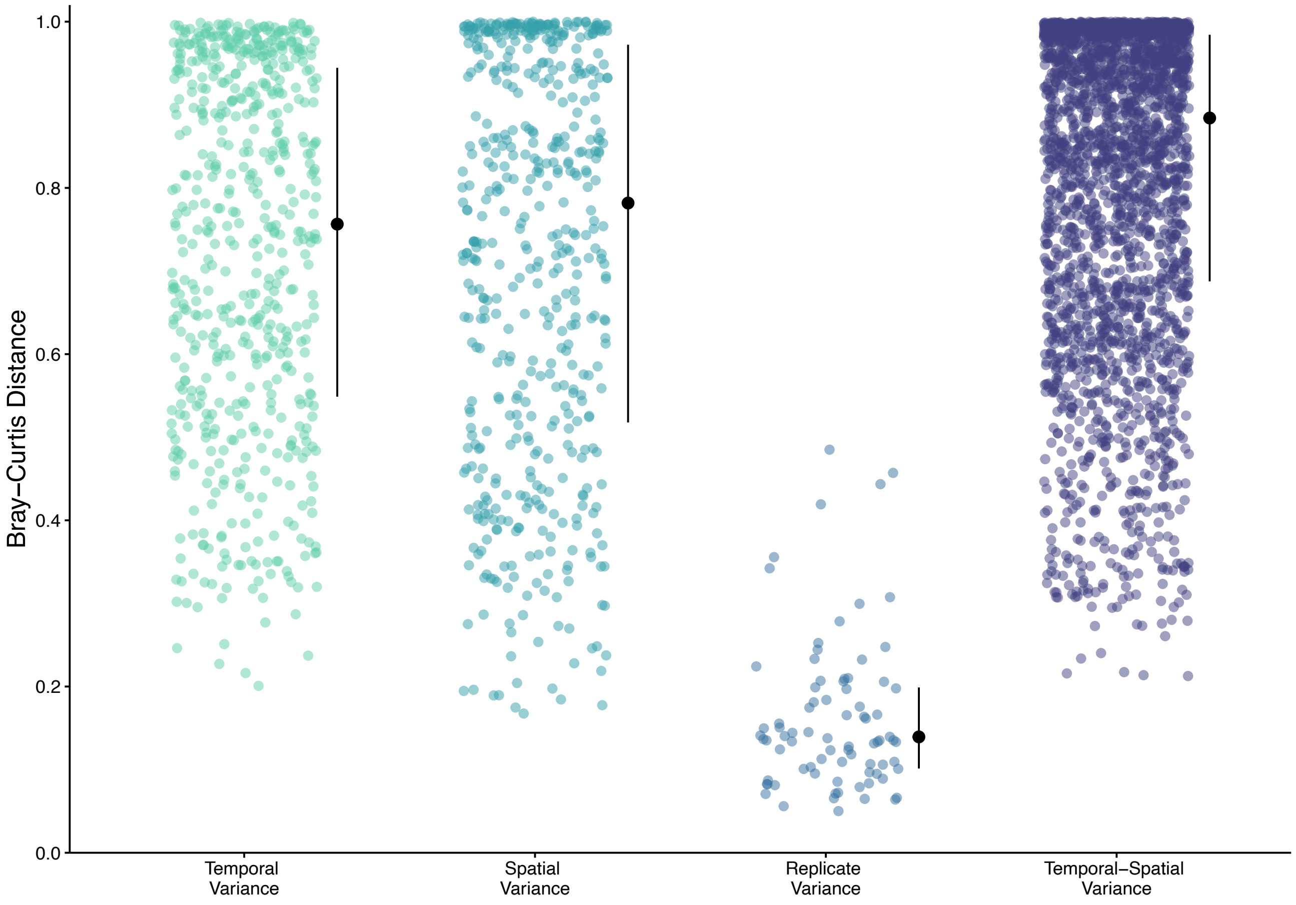


**Supplementary Figure 5.** Plot displaying the distribution of Bray-Curtis distances between samples. Colored dots represent ecological distances between samples. Black dots indicate the median, while black lines indicate the interquartile range (50% confidence interval).

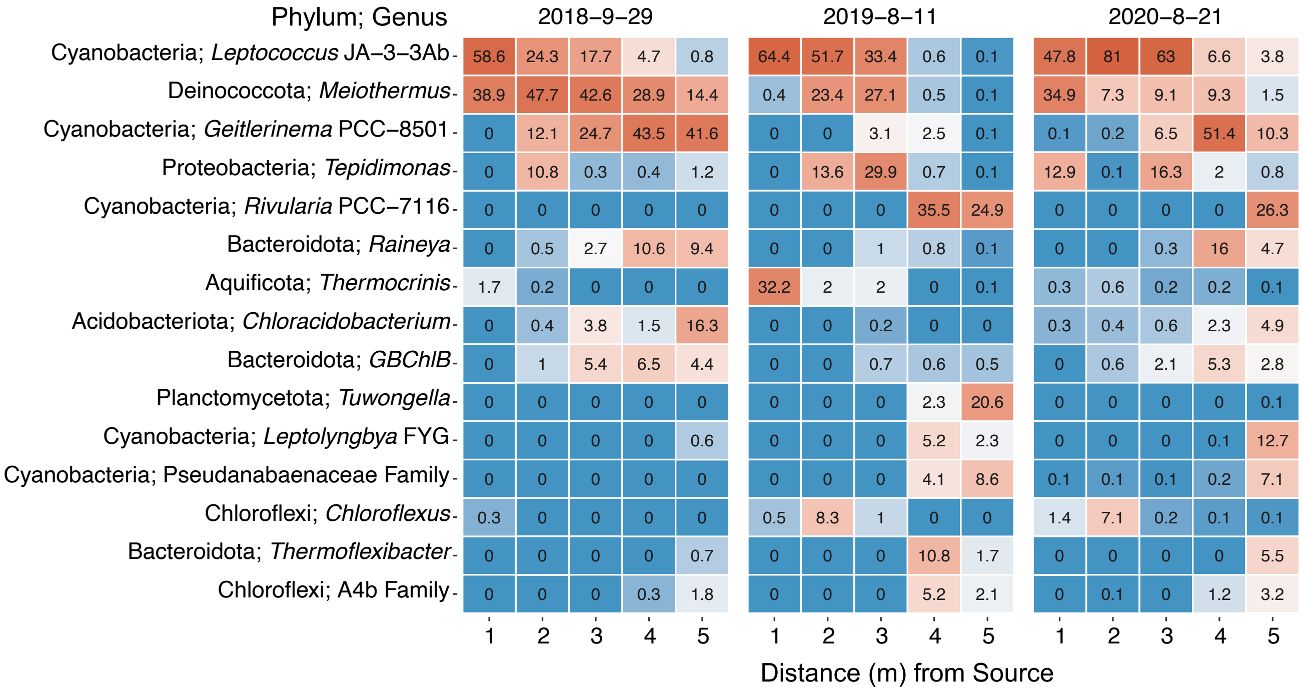


**Supplementary Figure 6.** Heat map of the top 15 ASVs within the bacterial and archaeal communities from 16S rRNA sequencing. ASVs are named by phyla and most likely genera. Values indicate the mean percent relative abundance. Data is faceted by sampling date and ordered down the sampling transect. Dates are shown as year-month-day.

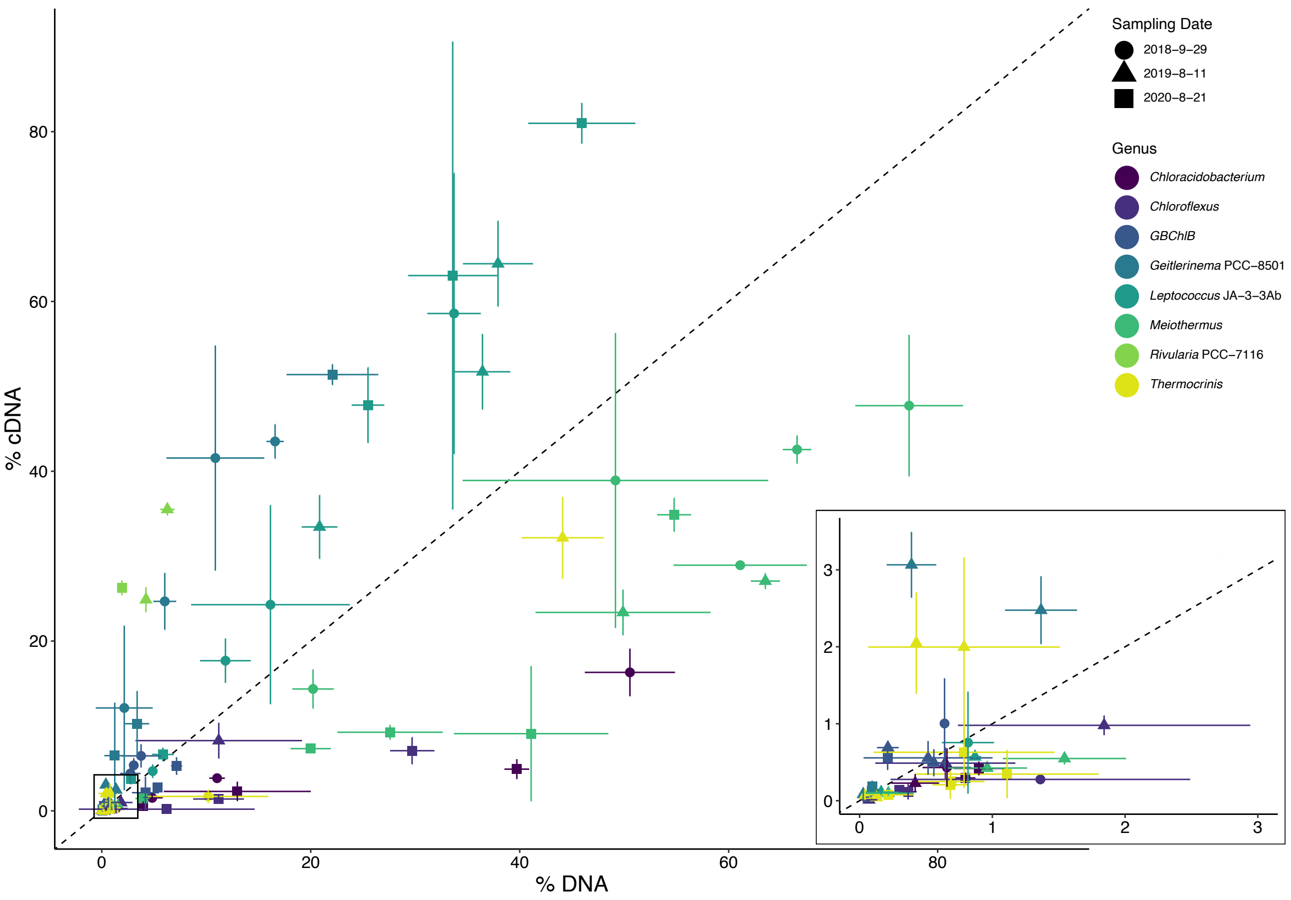


**Supplementary Figure 7.** Percentage of the top 8 ASVs as indicated by 16S rRNA sequence abundance. Inset of samples located within the black box of 0 to 3 % (c)DNA. Axes are percent relative abundance in either the cDNA (16S rRNA) or DNA (16S rRNA gene) sequencing data. Dots represent mean relative abundance and error bars indicate the standard deviation (n = 3) in the cDNA and DNA of biomass collected from the biofilms along the samples transect at Steep Cone Geyser. Genera classification is indicated by dot colors, while shapes indicate sampling date. Dotted line represents a 1:1 ratio of % cDNA to % DNA. Dates are shown as year-month-day.

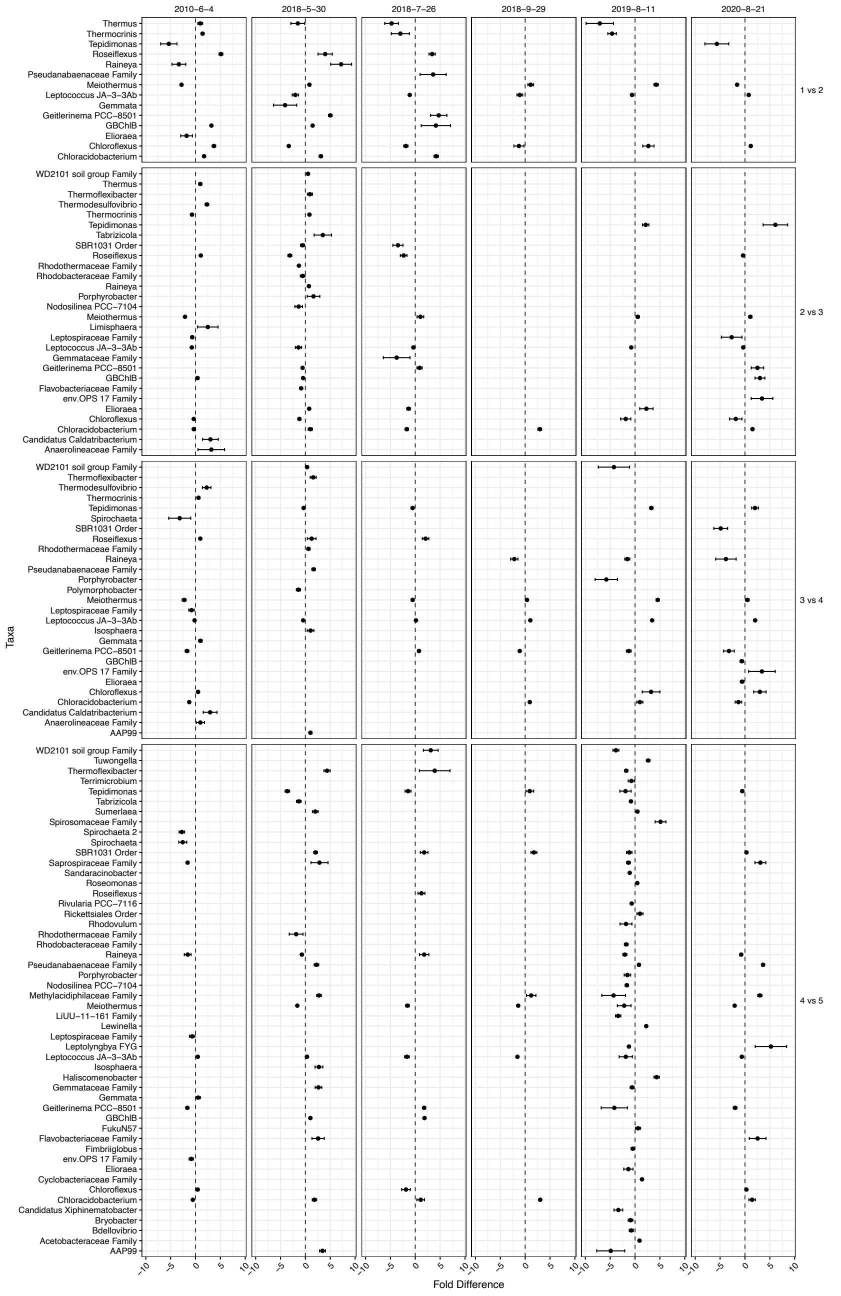


**Supplementary Figure 8.** Results from differential abundance analysis of 16S rRNA gene sequencing conducted using Corncob. Differential abundance comparison was conducted between adjacent sampling locations (right) for each sampling date (top). X-axis is $Log_2$ fold change. Listed taxa were determined to be statistically significant and are named by genera or best classification. All comparisons were performed sequentially. Therefore, the baseline comparison for each pairwise set is the lower number. Only taxa determined to be statistically significant for differential abundance after false discovery rate correction (*p-adj.* < 0.05) are shown. Positive values indicate the taxon increased in abundance from the baseline, while negative values indicate the taxon decreased in abundance from the baseline. Dates are shown as year-month-day.

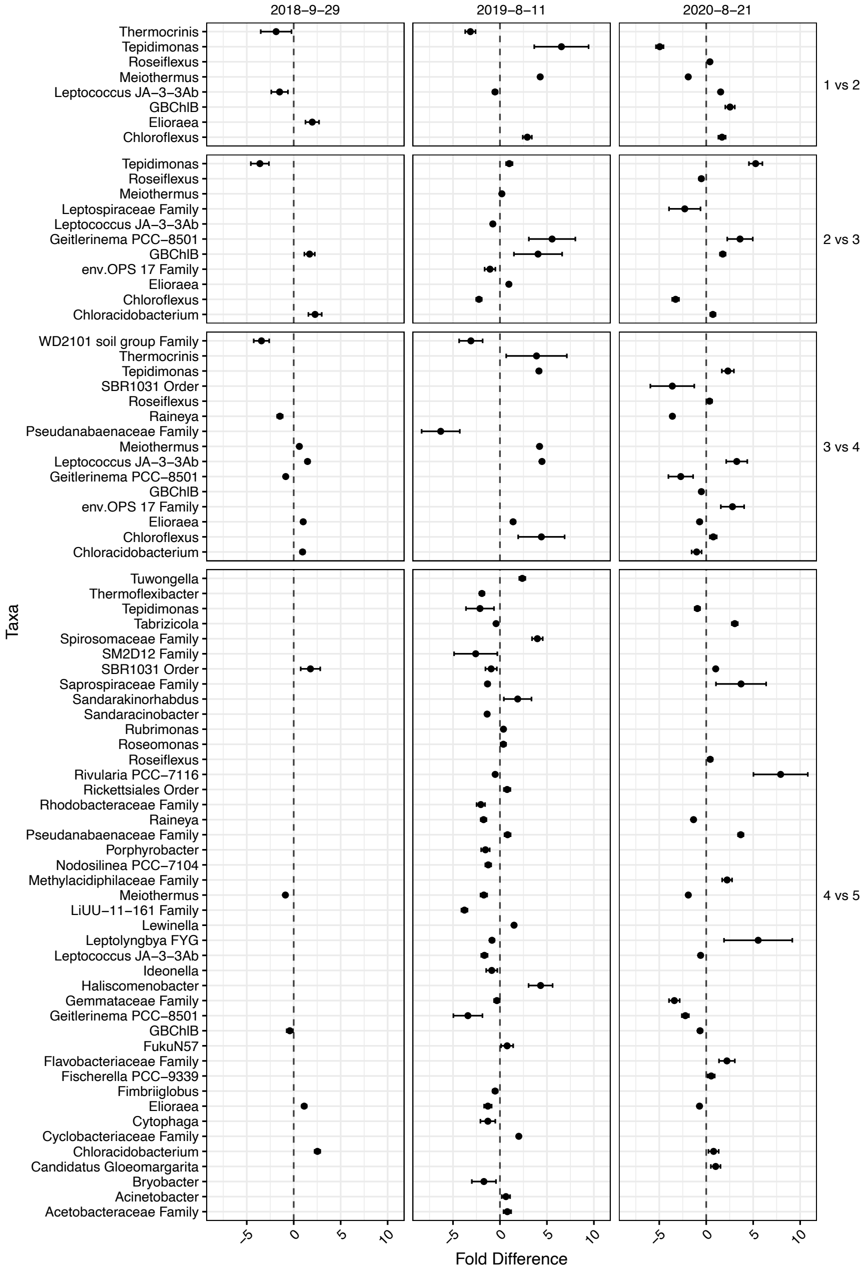


**Supplementary Figure 9.** Results from differential abundance analysis of 16S rRNA sequencing conducted using Corncob. Differential abundance comparison was conducted between adjacent sampling locations (right) for each sampling date (top). X-axis is $Log_2$ fold change. Listed taxa were determined to be statistically significant and are named by genera or best classification. All comparisons were performed sequentially. Therefore, the baseline comparison for each pairwise set is the lower number. Only taxa determined to be statistically significant for differential abundance after false discovery rate correction (*p-adj.* < 0.05) are shown. Positive values indicate the taxon increased in abundance from the baseline, while negative values indicate the taxon decreased in abundance from the baseline. Dates are shown as year-month-day.

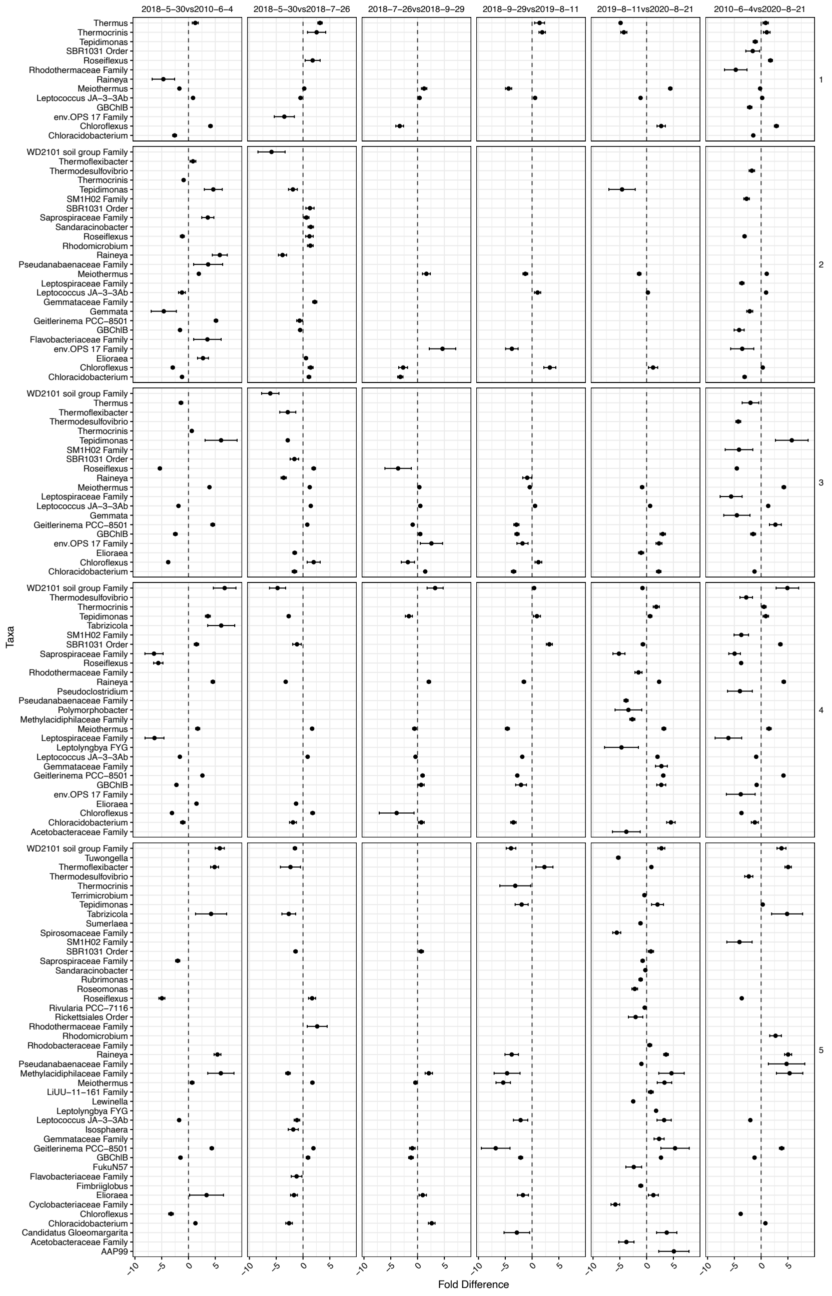


**Supplementary Figure 10.** Results from differential abundance analysis of 16S rRNA gene sequencing conducted using Corncob. Differential abundance comparison was conducted between sampling dates (top) for each sampling location (right). X-axis is $Log_2$ fold change. Listed taxa were determined to be statistically significant and are named by genera or best classification. All comparisons were performed sequentially. Therefore, the baseline comparison for each pairwise set is the lower number. Only taxa determined to be statistically significant for differential abundance after false discovery rate correction (*p-adj.* $< 0.05$) are shown. Positive values indicate the taxon increased in abundance from the baseline, while negative values indicate the taxon decreased in abundance from the baseline. Dates are shown as year-month-day.

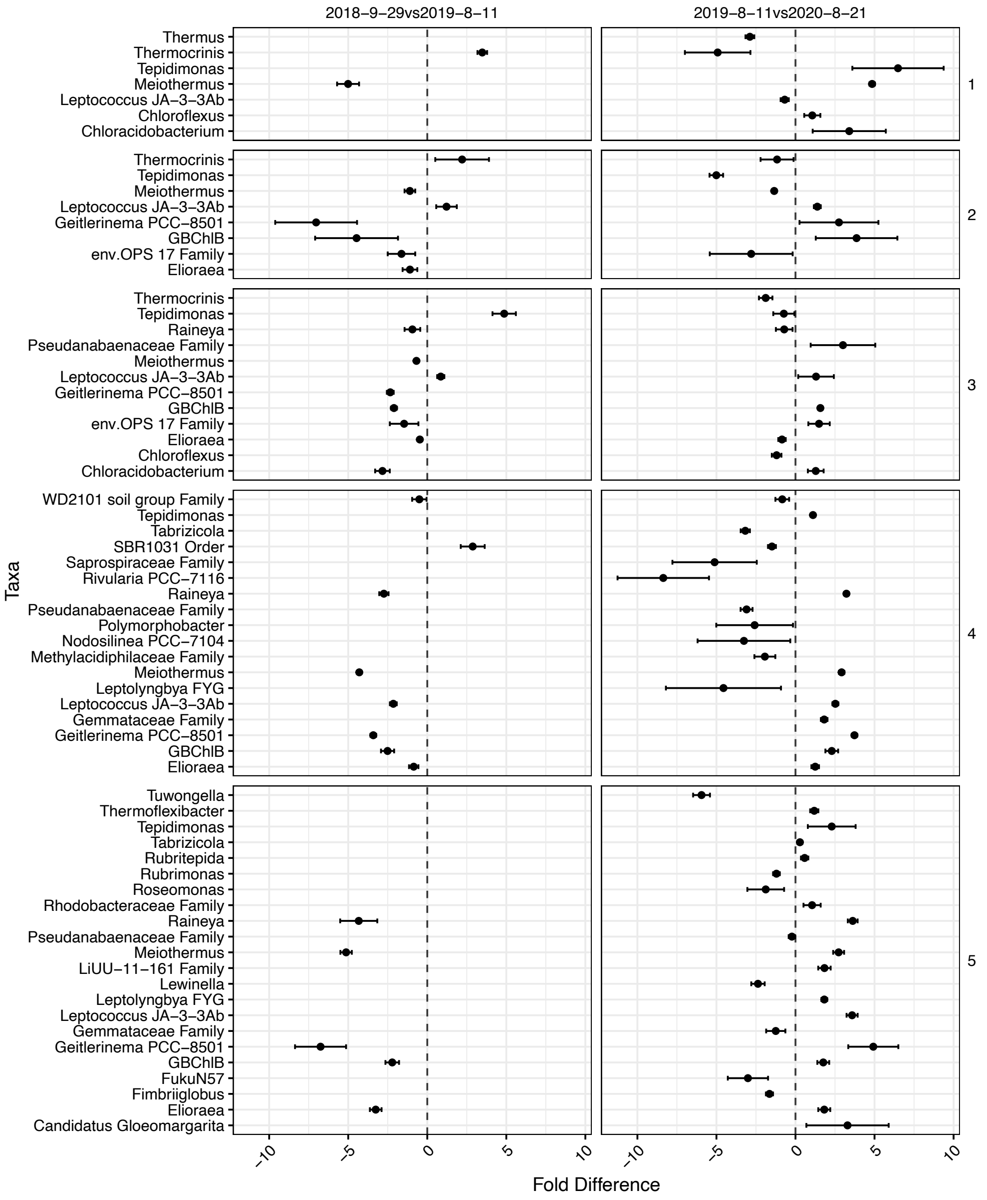


**Supplementary Figure 11.** Results from differential abundance analysis of 16S rRNA sequencing conducted using Corncob. Differential abundance comparison was conducted between sampling dates (top) for each sampling location (right). X-axis is $Log_2$ fold change. Listed taxa were determined to be statistically significant and are named by genera or best classification. All comparisons were performed sequentially. Therefore, the baseline comparison for each pairwise set is the lower number. Only taxa determined to be statistically significant for differential abundance after false discovery rate correction (*p-adj.* < 0.05) are shown. Positive values indicate the taxon increased in abundance from the baseline, while negative values indicate the taxon decreased in abundance from the baseline. Dates are shown as year-month-day.

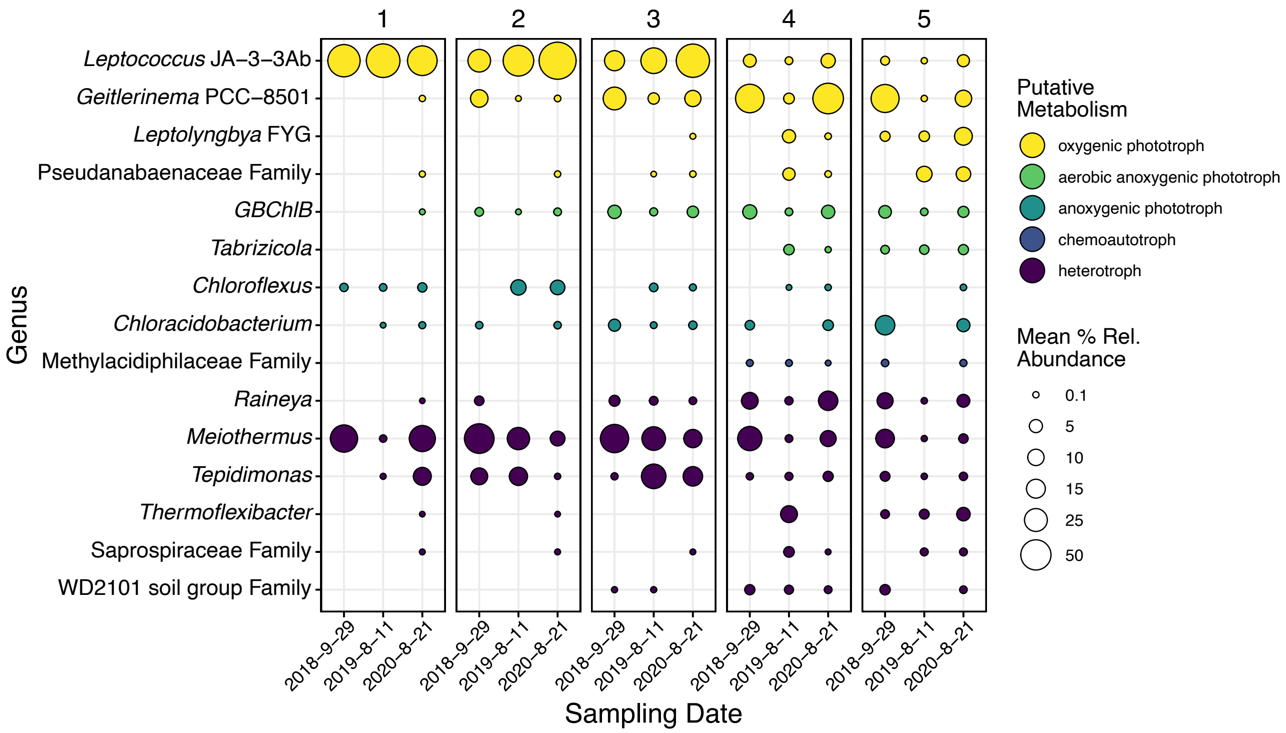


**Supplementary Figure 12.** 16S rRNA mean percent relative abundances of genera (or best classification) determined to be statistically significant by SIMPER, DNA differential abundance analysis, and cDNA differential abundance analysis. Panels are faceted by distance (m) from the hydrothermal source. Circle sizes indicate mean percent relative abundance. Colors indicate general putative metabolisms associated with the most abundant ASVs in each genus according to NCBI BLAST searches and literature review. Dates are shown as year-month-day.
